# SQSTM1/p62 accumulation is a hallmark of FLCN loss in Birt-Hogg-Dubé syndrome-associated kidney cancer

**DOI:** 10.64898/2026.01.14.699334

**Authors:** Halvor Ullern, Julie Aarmo Johannessen, Feyza Kasikci, Miriam Formica, Naghmeh Karimi Melve, Siri Andresen, Andreas Brech, Karol Axcrona, Kjersti Jørgensen, Lorant Farkas, Jorrit M. Enserink, Helene Knævelsrud

## Abstract

Birt-Hogg-Dubé syndrome (BHD) is an autosomal, dominant condition caused by *Folliculin* (FLCN) mutation and characterized by enhanced risk for kidney tumors. Previous studies have shown constitutive nuclear localization of the transcription factor TFEB and simultaneous hyperactivation of canonical MTORC1 signaling in the absence of FLCN. Here we assess the impact on autophagy under this situation of combined anabolic and catabolic activation. Using an established BHD patient-derived kidney cancer cell line, we confirmed that TFEB was permanently localized in the nucleus combined with an increase in canonical MTORC1 signaling, whereas bulk autophagy flux and LC3 lipidation was unaffected by FLCN status. However, we found that the autophagy receptor SQSTM1/p62 accumulated in enlarged puncta in the absence of FLCN. Finally, we recapitulate these findings in a Norwegian cohort of BHD kidney tumor samples. Our results demonstrate SQSTM1/p62 accumulation as a hallmark of FLCN loss, although SQSTM1/p62 appeared dispensable for anchorage-independent growth.

## Introduction

Birt-Hogg-Dubé (BHD) syndrome is an autosomal, dominant genetic disease characterized by fibrofolliculomas, lung cysts, pneumothorax and kidney tumors [1–3]. The most recent estimate of the penetrance of kidney tumors was 19-21% by the age of 70 in BHD patients with a male to female ratio of 2.5:1 [4]. Many BHD patients develop multiple and bilateral tumors, and the most common histological subtypes appear to be hybrid tumors followed by chromophobe renal cell carcinoma (RCC), oncocytomas and clear cell RCC [5,6]. There is currently no specific kidney cancer treatment for BHD-patients besides surgery when the tumor has reached a certain size and standard treatment according to guidelines for sporadic RCC in the case of metastatic disease [7–9].

BHD is caused by loss-of-function germline mutations in the gene that encodes folliculin (FLCN). *FLCN* is considered a tumor suppressor gene since its loss may lead to kidney tumors. Normal FLCN is a 64-kDa protein that is conserved across species, but shares no homology to any other known proteins [10,11]. FLCN forms heterodimers with folliculin-interacting protein 1 (FNIP1) or FNIP2 [12,13]. Under physiological conditions, the FLCN/FNIP heterodimer acts as an amino acid sensor at the lysosome. When amino acids are sparse, FLCN is inactive as a part of the lysosomal folliculin complex (LFC) [14], and it becomes activated when amino acids are abundant [15]. Active FLCN activates the kinase mechanistic Target of Rapamycin (MTOR) through its GTPase Activating protein (GAP) activity towards Ras related GTP-binding protein (RAG) C/D [16,17].

MTOR is a conserved serine/threonine protein kinase that is part of at least two protein complexes: MTOR complex 1 and 2 (MTORC1 and MTORC2, respectively). MTORC1 combines information from environmental cues like growth factors, nutrient status and energy levels and thereby governs cell growth and proliferation. MTORC1 activation depends on two sets of GTPases and their nucleotide state: Ras homolog enriched in brain (RHEB) and the RAG GTPases. RAGs C or D, which are active when GDP-bound, form a heterodimer with either RAG A or B, which are active when GTP-bound [18]. The RAGs facilitate the recruitment of MTORC1 to the lysosomal surface where it can be activated when nutrients are abundant [19,20].

Phosphorylation targets of MTORC1 encompass RPS6K and EIF4EBP1/4E-BP1, involved in protein synthesis, along with ULK1 and ATG13, pivotal for macroautophagy/autophagy, as well as the microphtalmia (MiT)-TFE transcription factors, especially Transcription Factor EB (TFEB) and E3 (TFE3). Under nutrient sparse conditions, MTORC1 no longer exerts inhibitory phosphorylation of ULK1, ATG13 and TFEB/TFE3, triggering the initiation of autophagy [18]. The exact relationship between FLCN loss and the activity of the canonical MTORC1 targets is not fully understood, indeed both increased and decreased activity has been reported [21–26].

TFEB and TFE3 drive transcription of genes related to autophagy and lysosomal biogenesis [27–29]. Loss of FLCN results in constitutive de-phosphorylation and subsequent nuclear localization of TFEB and TFE3 [16,30]. The phosphorylation of TFEB and TFE3 is mediated by amino acid-dependent activation of RAG C/D through FLCN, but not by growth factor dependent RHEB activity, like other substrates of MTORC1 including RPS6K and EIF4EBP1/4E-BP1 [14,21,31,32]. The lysosomal MTORC1-TFEB-RAG-Ragulator megacomplex has been determined with cryo-EM, showing that TFEB phosphorylation by MTORC1 is regulated through a mechanism dependent on RAGC^GDP^ [33]. Conversely, TFEB or TFE3 may be able to exert a positive regulation on MTORC1 and its canonical targets through transcriptional activation of RAG C/D [34]. TFEB and TFE3 have also been shown to drive kidney cystogenesis and tumorigenesis [35].

Although substantial progress has been made in the last years elucidating FLCN’s role in MTOR-, AMPK- and TFEB signaling, much is still unknown about how FLCN affects autophagy, a catabolic process in which autophagosomes sequester portions of the cytoplasm before fusing with lysosomes for degradation of the encapsulated material [36]. Some studies have indicated that FLCN loss may lead to decreased autophagic flux, accumulation of SQSTM1/p62 and dysregulation of LC3 and GABARAP family proteins [25,37–39]. FLCN loss is reported to both increase the activity of MTORC1, which promotes anabolism and suppresses autophagy, and to increase the nuclear localization of TFEB/TFE3, which promotes catabolism through transcription of autophagic and lysosomal genes. To address this conundrum, we aimed to identify the impact of FLCN loss on autophagy in BHD syndrome.

## Results

### TFEB is Constantly Nuclear Localized in the Absence of FLCN

To delineate FLCN-dependent effects on autophagy, we utilized the established BHD-patient derived FLCN-deficient cell line (UOK257) [40] and compared it to the same cell line where FLCN expression had been restored by stable expression (UOK257-2) [13]. Several studies have shown that the loss of FLCN results in the constitutive nuclear localization and hyperactivation of both TFEB [16,35,41,42] and TFE3 [30,32,42,43]. Consistently, we found that TFEB remained constitutively localized to the nucleus in FLCN-deficient UOK257 cells regardless of the conditions, including starvation, refeeding, or Torin 1 treatment (Fig. 1A). In contrast, FLCN-reconstituted UOK257 cells exhibited cytosolic localization of TFEB under fed conditions, with nuclear translocation occurring upon starvation and subsequent return to the cytosol after nutrient replenishment (Fig. 1A-B). Notably, TFEB was retained in the nucleus when MTOR was inhibited by Torin 1 during nutrient replenishment. These observations were validated through nuclear/cytosolic fractionation immunoblot experiments (Fig. 1C-E). Correspondingly, phosphorylation of TFEB at Ser11 occurred only in the presence of FLCN, with the phosphorylated form found in the cytosolic fraction – and only in the absence of MTOR inhibition by Torin 1 – indicating a dependency on MTOR activity (Fig. 1C-E). Collectively, these findings underscore that in the FLCN-deficient UOK257 cell line, TFEB is persistently localized in the nucleus.

**Figure 1.**
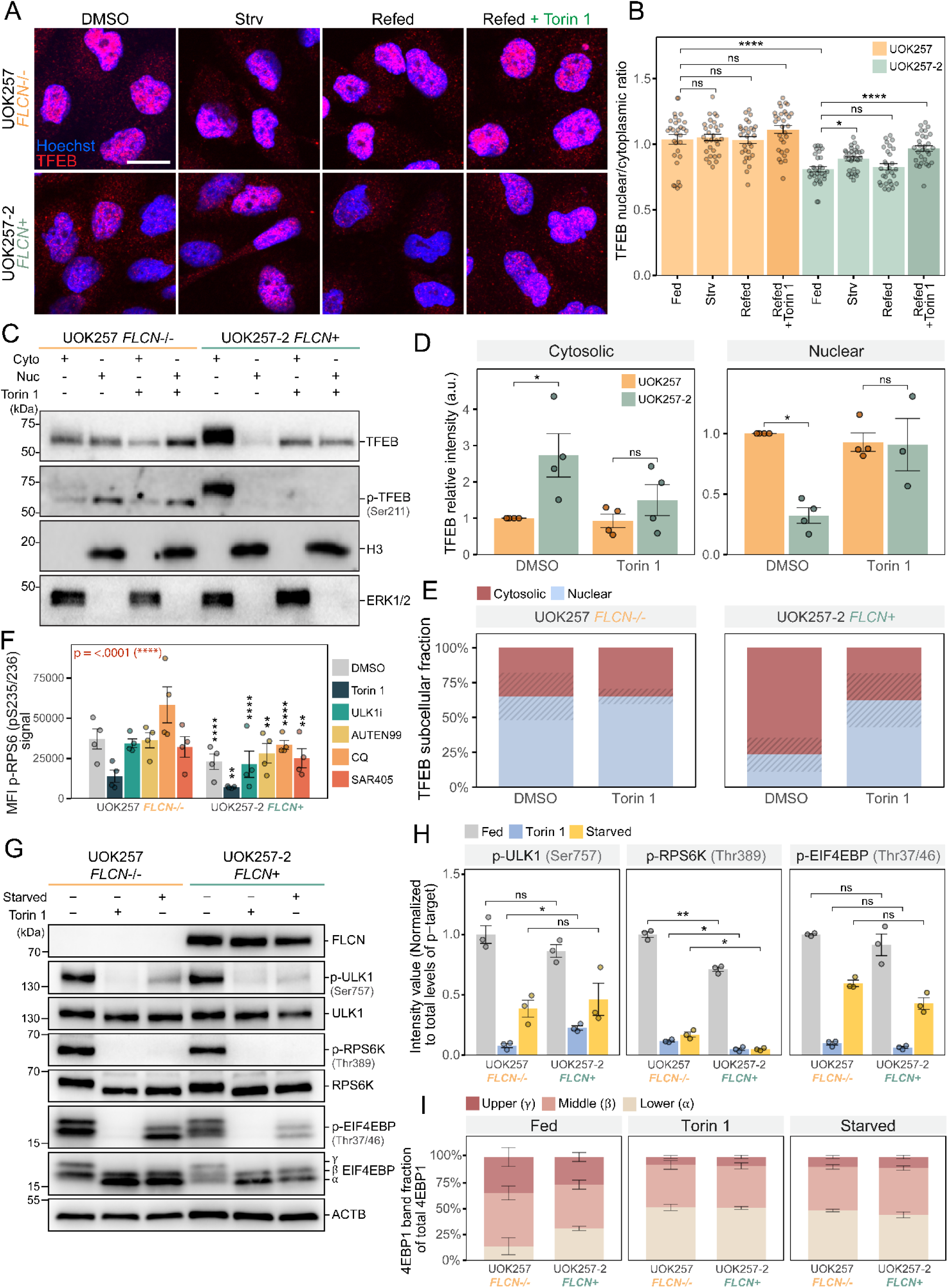
FLCN reconstitution in UOK257 cells increases TFEB phosphorylation and reduces canonical MTOR signaling. (**A**) Immunofluorescence analysis showing subcellular localization of TFEB (red) in either fed (DMSO), starved (Strv), starved and refed (Refed) or starved and refed cells with Torin1 treatment during refeeding. Scale bar: 20 µM. (**B**) Quantification of images in panel A, displayed as nuclear/cytoplasmic ratio of TFEB intensity. Bar plots show mean ± SEM (N=10 images in 3 biological replicates). Statistical analysis was performed using two-sided t-test followed my Benjamini-Hochberg’s correction for multiple comparisons. (**C**) Nuclear/cytosol fractionation western blot showing subcellular localization of TFEB. Histone 3 (H3) and ERK1/2 were used as control for the nuclear and cytoplasmic fraction, respectively. (**D**) Quantification of cytosolic and nuclear protein levels from panel C. Bar plots show mean ± SEM (N=3-4), normalized to UOK257 *FLCN-/-* DMSO within each experiment. Statistical analysis was performed using Wilcoxon’s Rank Sum test. (**E**) Data from panel D, displayed as stacked bar plot to showcase the distribution of cytosolic (red) versus nuclear (blue) fractions within UOK257 cell lines. Hatched area is ± SD of fraction distribution (N=3-4 replicates). (**F**) Median fluorescent intensity (MFI) values of phospho-RPS6 (p-Ser235/Ser236) signal measured with phospho flow cytometry. P-value on plot indicate results of one-way ANOVA test of overall treatment-specific response in-between cell lines. P-values above bars in UOK257-2 indicate statistical significance compared to the respective treatment in UOK257, as determined by Tukey’s multiple comparisons test. Bar plots show mean ± SEM (N=4 replicates). (**G**) Western blot of canonical MTORC1 phosphorylation substrates in either fed, starved or Torin 1 treated UOK257 cells. In EIF4EBP blot, γ, β and α annotates three distinct bands of different weight. (**H**) Quantification of phosphorylated MTORC1 substrates from panel G, comparing fed samples (gray) to Torin 1 treated (blue) or starved samples (yellow). Values are relative to total level of the respective phospho-substrate and normalized to UOK257 *FLCN*-/- control. For p-EIF4EBP, all bands shown in immunoblot in G was used for the quantification. Bar plots show mean ± SEM (N=3 replicates). Statistical analysis was performed using one-way ANOVA with Tukey’s multiple comparisons test. (**I**) Quantification of EIF4EBP protein levels from panel G, shown as fractions of bands with different weight relative to total EIF4EBP levels in fed, Torin 1 treated and starved samples. Error bars are ± SD (N=3 replicates). ns: not significant (*P* > 0.05); \**P* < 0.05, \*\**P* < 0.01, \*\*\**P* < 0.001, \*\*\*\**P* < 0.0001

### Canonical MTORC1 activity promoting anabolism is upregulated in the absence of FLCN

Given that TFEB serves as a key regulator of autophagy-related genes, we hypothesized that autophagy-related pathways would be dysregulated in FLCN-deficient UOK257 cells. To elucidate which autophagy-related pathways are differentially regulated in UOK257 (*FLCN* -/-) compared to UOK257-2 (FLCN reconstituted), cells were treated with various pharmacological agents that are known to affect autophagy: The MTOR inhibitor Torin 1 which induces autophagy [44]; the MTMR14 inhibitor AUTEN99, which increases autophagosome biogenesis [45]; the ULK1 inhibitor MRT68921 which prevents the initiation of autophagy [46]; the PIK3C3 inhibitor SAR405, which impedes autophagosome biogenesis [47]; and Chloroquine, which impairs lysosomal acidification and autophagosome and lysosome fusion [48]. Phospho-flow cytometry analysis was subsequently performed using 13 distinct antibodies targeting key signaling events linked to receptor tyrosine kinase signaling, MTOR signaling, proliferation, autophagy, cellular stress, and apoptosis pathways. The vehicle-treated (DMSO) cells showed mostly similar patterns of phosphorylation levels in the presence or absence of FLCN (Fig. S1A). Notably, the only significant finding across all tested conditions was the downregulation of RPS6 phosphorylation (pS235/236) in the FLCN-restored cell line (Fig. 1F). Although increased phosphorylation of p44/42 MAPK (pT202/Y204) was also noted, it did not reach statistical significance under these conditions (Fig. S1A).

As RPS6 phosphorylation on pS235/236 was increased in FLCN-deficient UOK257 across all conditions, we investigated MTORC1 targets further. The precise effect on the canonical MTOR targets upon loss of FLCN is still not fully resolved [21–26], thus, we aimed to determine their phosphorylation levels in the UOK257 cell lines (Fig. 1G). Cells were either fed, starved, or treated with Torin 1. Upon Torin 1 treatment, the expected reduction of phosphorylation of canonical MTOR targets ULK1 (pSer757), RPS6K (pThr389) and EIF4EBP1 (pThr37/46) was observed in both UOK257 and UOK257-2. In basal fed conditions, p-RPS6K levels were reduced in UOK257-2 compared to UOK257, and the same pattern was observed for starved cells (Fig. 1G-H). In the case of EIF4EBP1, in fed cells lacking FLCN, EIF4EBP1 existed predominantly as the slower migrating upper two bands (γ, β), whereas in FLCN-reconstituted cells a shift towards the faster migrating lower band (α) was observed, indicating reduced phosphorylation (Fig. 1I). As expected, both cell lines exhibited a shift towards the lower EIF4EBP bands upon Torin 1 treatment or starvation, consistent with reduced phosphorylation (Fig. 1I). However, ULK1 did not show significantly reduced phosphorylation at Ser757 in the FLCN-negative UOK257 cells. These findings indicate that the MTOR-mediated anabolic signaling through RPS6K and EIF4EBP1 is increased in this cell model of FLCN deficiency, without significant effects on the inhibitory phosphorylation on ULK1 Ser757.

### Autophagy flux in UOK257 cells does not depend on FLCN

Given the observed dysregulation of MTOR and TFEB in FLCN-deficient cells, we hypothesized that this aberrant signaling would result in increased autophagic activity. Therefore, we subsequently examined autophagy flux using several assays. The ATG8 protein family in mammals includes three constituents each from the microtubule-associated proteins 1A/1B light chain 3 (MAP1LC3) family (LC3A, LC3B, LC3C) and gamma-aminobutyric acid type A receptor-associated protein family (GABARAP, GABARAPL1, GABARAPL2). Taking LC3 as an example of ATG8 family proteins, upon induction of autophagy, cytosolic LC3 (LC3-I) is conjugated to phosphatidylethanolamine (PE), creating the lipidated form (LC3-II), which is found in the membrane of autophagic vesicles, and is later degraded by lysosomal hydrolases upon fusion of autophagosomes with autolysosomes. LC3-II is thus used as a marker for autophagy and LC3 flux can be assessed by adding a lysosomal inhibitor to measure the full amount of LC3 lipidation during the treatment [49].

First, the LC3 flux was assessed by measuring lipidation of LC3 with immunoblotting after treatment with vacuolar H+ ATPase (V-ATPase) inhibitor Bafilomycin A1 (BafA1). This revealed comparable levels of basal LC3 flux in fed conditions for both cell lines, as seen by equal levels of LC3-II accumulation after BafA1 treatment. Moreover, both cell lines displayed the expected increase in LC3 flux upon nutrient deprivation, indicating capable autophagic machinery (Fig. 2A-B). The observed accumulation of LC3-II in the presence of BafA1 was inhibited by SAR405, which suggested a dependency on canonical PIK3C3 activity in both cell lines. On the transcriptional level, RT-qPCR for the different members of the LC3 family showed that LC3A, LC3B, GABARAP and GABARAPL2 mRNA levels were similar between the two cell lines, whereas LC3C expression was approximately 15-fold higher and GABARAPL1 levels were marginally lower in UOK257-2 (Fig. S2A).

**Figure 2.**
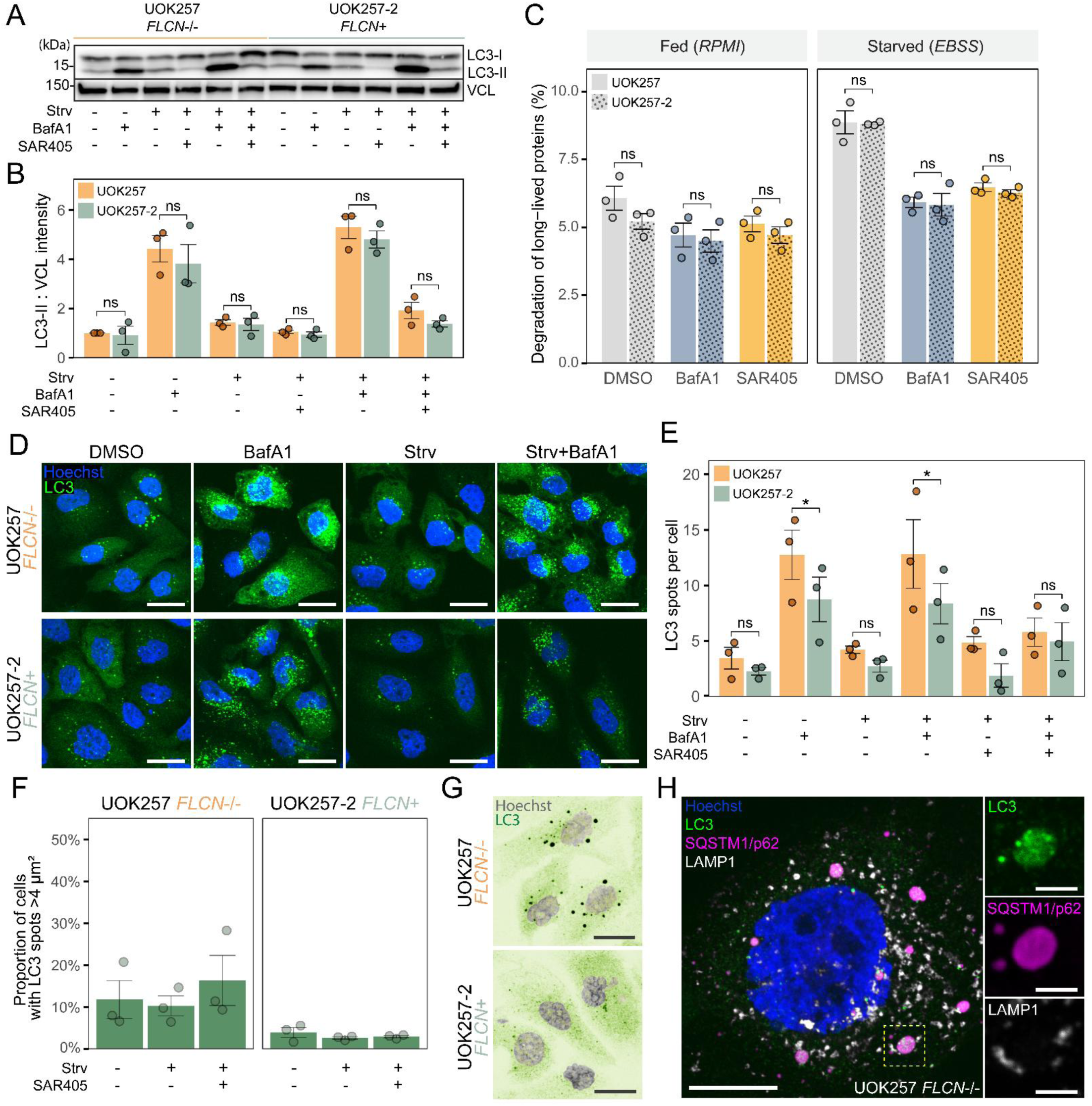
Autophagy flux in UOK257 cells is unaffected by FLCN reconstitution. (**A**) Western blot analysis of LC3 protein levels and LC3 lipidation in UOK257 cells. (**B**) Quantification of lipidated LC3 levels from western blot in panel A. LC3-II levels are shown relative to VCL/vinculin and normalized to UOK257 *FLCN-/-* DMSO control. Bar plots show mean ± SEM (N=3 replicates). Statistical analysis was performed using one-way ANOVA with Tukey’s multiple comparisons test. (**C**) Quantification of long-lived protein degradation (LLPD) assay showing percentage of degradation of long-lived proteins for fed (RPMI) and starved (EBSS) samples. Bar plots show mean ± SEM (N=3). Statistical analysis was performed using one-way ANOVA with Tukey’s multiple comparisons test. (**D**) Immunofluorescence analysis of LC3 (green) in fed (DMSO) or starved (Strv) UOK257 cells, with or without Bafilomycin A1 (BafA1) treatment. Scale bar: 25 µm. (**E**) Quantification of LC3 spots per cell from images in panel D. Bar plot shows mean ± SEM (N=3 replicates). Statistical analysis was performed using one-way ANOVA with Tukey’s multiple comparisons test. (**F**) Quantification of proportion of UOK257 cells with LC3 puncta > 4 µm^2^. Bar plots show mean ± SEM (N=3 replicates). (**G**) Representative images of LC3 puncta (green) in starved UOK257 cells treated with SAR405. Nuclei are shown in gray. Scalebar: 25 µm. (**H**) Immunofluorescence analysis of colocalization of LC3 (green), SQSTM1/p62 (magenta) and LAMP1 (gray). Representative image of UOK257 cell under basal fed conditions. Scale bars: merge panel: 10 µm; inserts: 2 µm. ns: not significant (*P* > 0.05); \**P* < 0.05, \*\**P* < 0.01, \*\*\**P* < 0.001, \*\*\*\**P* < 0.0001.

To quantify autophagic flux, the long-lived protein degradation (LLPD) assay was employed [50]. Cells were cultured in the presence of a radiolabeled amino acid (valine ^14^C) to label the proteome. Short-lived proteins were then chased away to follow the rate of degradation of long-lived proteins, and degradation rates were compared across fed and starved states. Moreover, cells were treated with BafA1 and SAR405 to inhibit the autophagy pathway at the level of lysosomal degradation or PIK3C3 activity, respectively. In agreement with LC3 immunoblotting, the results of LLPD indicated that the basal autophagic flux was similar between the cell lines and that both cell lines induced autophagy to a similar extent as a response to starvation (Fig. 2C).

To complement these results with visualization of autophagosome formation and turnover, we employed immunofluorescence analysis of LC3. This revealed similar levels of LC3 puncta in fed cells, and no differences were observed between the cell lines when inducing autophagy with starvation (Fig. 2D-E). Aligning with immunoblotting results, BafA1 treatment resulted in the accumulation of LC3 positive vesicles in both cell lines, although to significantly higher levels in cells lacking FLCN (UOK257 cells, Fig. 2D-E). This finding could indicate a slightly elevated LC3 flux in UOK257. Interestingly, we observed that a proportion of the LC3 spots formed in FLCN-lacking UOK257 cells were morphologically distinct from those in UOK257-2 cells, with large LC3 spots observed in the absence of FLCN (Fig. 2F). Notably, these large puncta persisted upon inducing autophagy by starvation and were not resolved following PIK3C3 inhibition by SAR405 (Fig. 2G). To determine the entity of the enlarged LC3-positive structures, we assessed colocalization with the autophagy cargo receptor SQSTM1/p62 (sequestosome 1) [51]. Immunofluorescence analysis demonstrated that the enlarged structures in UOK257 cells were densely positive for SQSTM1/p62, while LC3 signal associated sparsely as discrete puncta. Moreover, the structures were uniformly negative for lysosomal membrane LAMP1 (Fig. 2H).

Collectively, these findings suggest that FLCN loss and subsequent constitutive TFEB nuclear localization does not result in a permanently increased autophagy flux.

### SQSTM1/p62 accumulates in FLCN-negative UOK257

Given the identification of SQSTM1/p62-dense, LAMP1 negative structures, SQSTM1/p62 expression and turnover were profiled in UOK257 cells. SQSTM1/p62 directly interacts with the ATG8 proteins and is primarily degraded by the autophagy pathway. SQSTM1/p62 may accumulate through reduced or impaired degradation in the autolysosome and is therefore often used as a marker of autophagic flux [49,52,53]. Immunoblotting for SQSTM1/p62 on lysates from cells under the same conditions as used to investigate LC3 flux revealed an accumulation of SQSTM1/p62 in UOK257 across all conditions (Fig. 3A-B), although starvation did induce a decline in total SQSTM1/p62 levels, which was blocked by the presence of BafA1. This indicated that at least part of the SQSTM1/p62 pool was turned over by autophagy also in the absence of FLCN. Increased mRNA levels of SQSTM1/p62 were measured in UOK257 (Fig. 3C), while levels of another autophagy receptor, NBR1, exhibited a decreasing trend (Fig. S3A). Consistently, immunofluorescence analysis confirmed elevated levels of SQSTM1/p62 in UOK257 (Fig. 3D-F). Notably, cytosolic SQSTM1/p62-positive aggregates larger than 6 µm^2^ were detected in UOK257 but were rare in FLCN-restored UOK257-2 (Fig. 3G). These spots accumulated when PIK3C3 activity was blocked by SAR405 (Fig. 3G). SQSTM1/p62 is known to be found in both membrane-free cytoplasmic bodies, as well as autophagosomes [52]. Ultrastructural analysis by electron microscopy (EM) showed that the SQSTM1/p62 positive structures observed in UOK257 were large membrane-free condensates of varying sizes (Fig. 3H).

**Figure 3.**
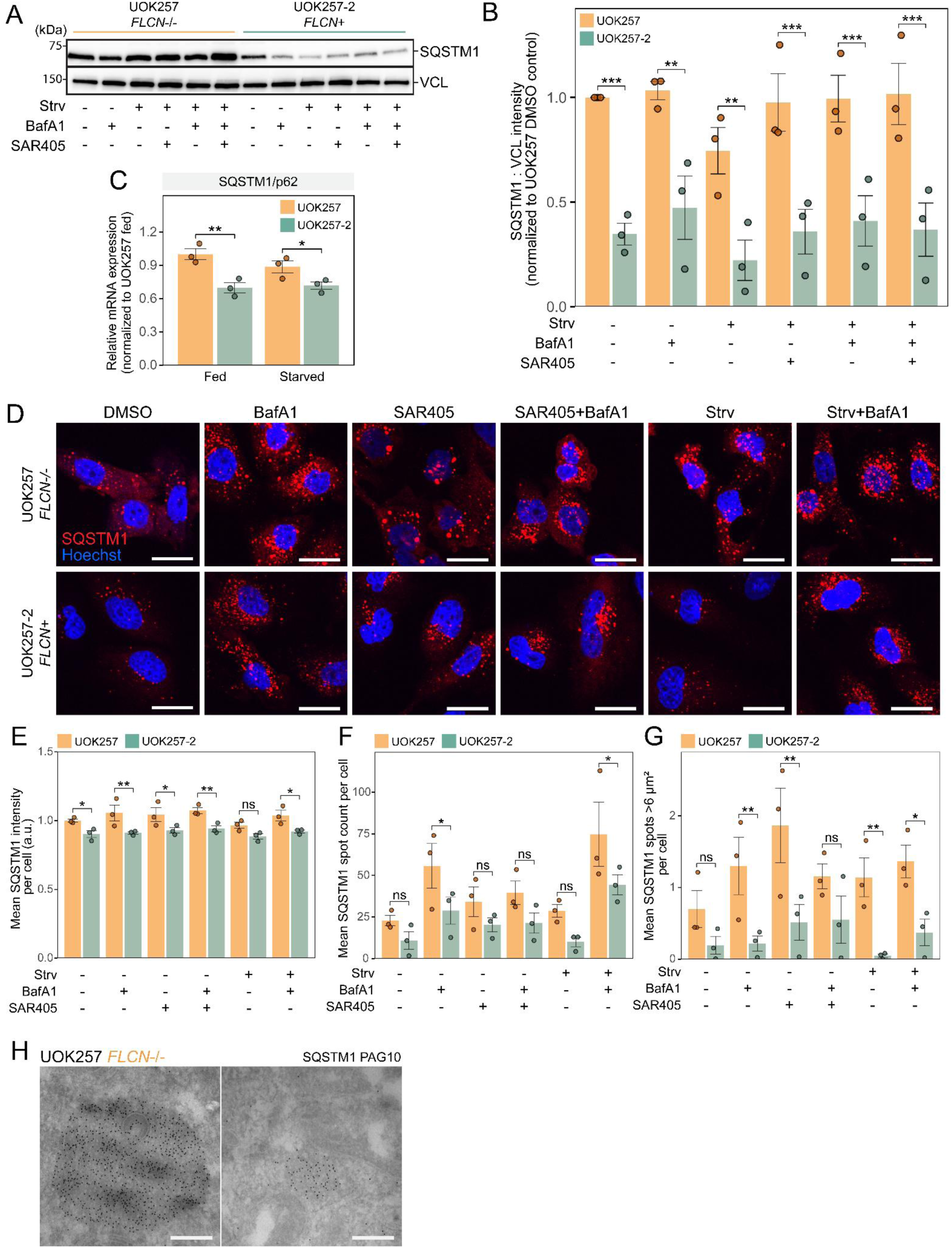
SQSMT1/p62 accumulation in large foci are characteristic features of UOK257 cells deficient in FLCN. (**A**) Western blot analysis of SQSTM1/p62 protein levels in UOK257 cells. (**B**) Graphs quantifying the western blots from A. Bar plots show mean ± SEM (N=3). Statistical analysis was performed using one-way ANOVA with Tukey’s multiple comparisons test. (**C**) Relative mRNA expression levels of *SQSTM1* measured by qRT-PCR in UOK257 cells. Expression levels are normalized to UOK257 *FLCN-/-* fed sample. Statistical analysis was performed using one-way ANOVA with Tukey’s multiple comparisons test. (**D**) Immunofluorescence analysis of SQSTM1/p62 (red) in UOK257 cells. Scale bar: 25 µm. (**E-G**) Quantification of mean SQSTM1/p62 intensity per cell (E), total SQSTM1/p62 spots (F) and number of large SQSTM1/p62 spots (> 6 µm^2^) (G) from images in panel D. Bar plots show mean ± SEM (N=3). Statistical analysis was performed using one-way ANOVA with Tukey’s multiple comparisons test. (**H**) Immuno-EM of SQSTM1/p62 in *FLCN*-/- cells show very dense labeling on cytosolic structures that are not surrounded by any limiting single membrane or autophagic membrane. The size of these structures varies considerably from 200 nm up to 2 µm. Labeling was performed with p62-antibody followed by 10 nm ProteinA-gold (PAG 10). Scale bar: 500 nm.

Taken together, we found that the FLCN-deficient cell line accumulated intracellular SQSTM1/p62-positive aggregates. This accumulation is not a result of decreased autophagic flux, but may be explained by an increased SQSTM1/p62 transcription.

### SQSTM1/p62 also accumulates in kidney tumors from BHD patients

To explore the clinical relevance of these findings, we investigated kidney tumors along with adjacent normal tissue from eight unrelated BHD syndrome patients (Table 1 and further histological details in Table S1). Our cohort showed a variety of histological tumor subtypes consistent with previous reports [6] (Fig. S4A, Table 1 and Table S1). To analyze levels of autophagy-related proteins specifically in the tumor cells of the BHD patient samples, we trained an object classifier in QuPath to distinguish tumor cells from stromal cells based on morphological appearance [54]. The classifier was validated by a trained pathologist. In accordance with our results from the UOK257 cell lines, we found TFEB in the nucleus of tumor cells in the BHD patient samples, while stromal cells were negative. (Fig 4A-B, Fig S4B).

**Figure 4:**
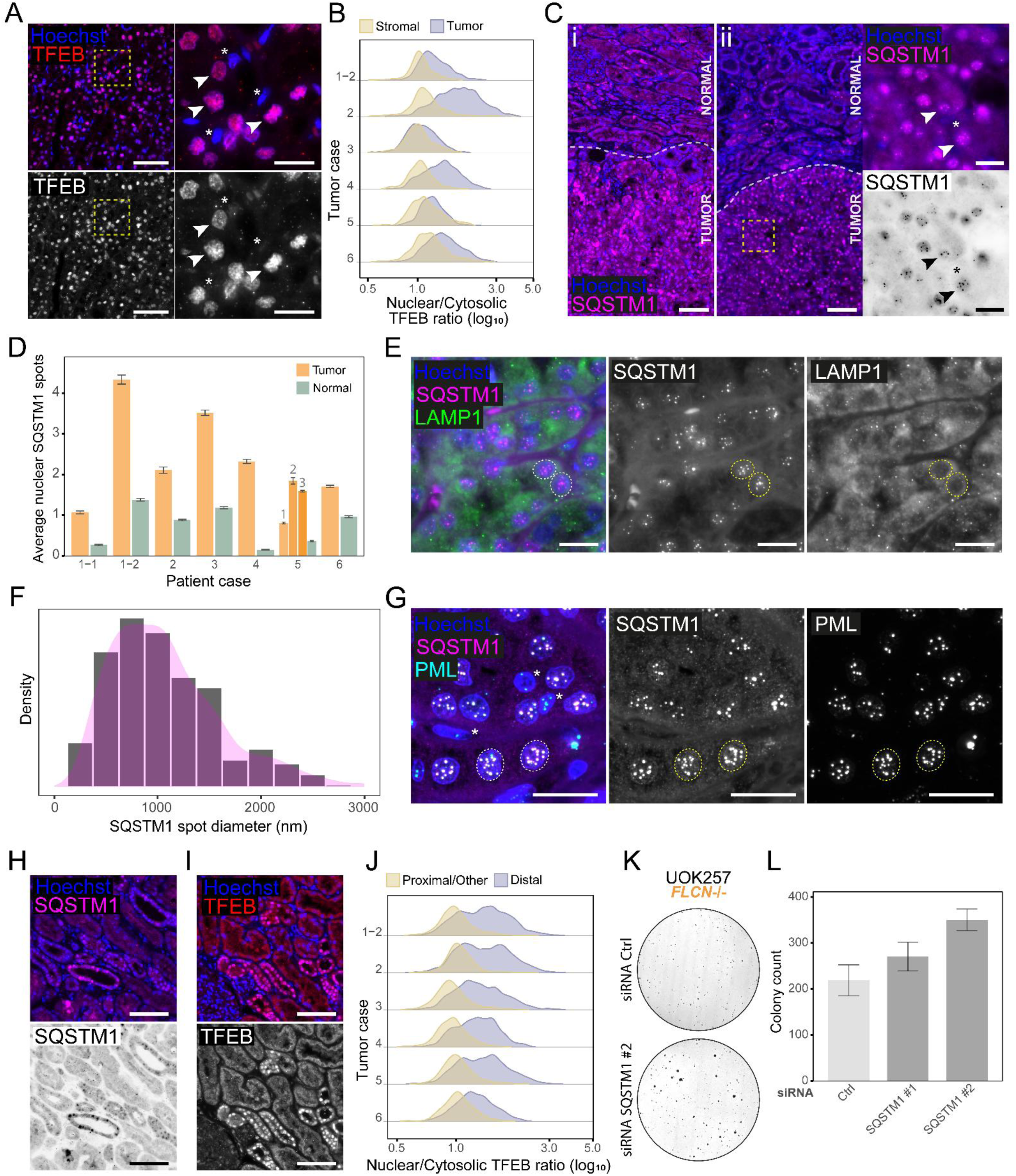
Nuclear SQSTM1/p62 colocalizing with PML bodies is a hallmark of BHD tumors. (**A**) Representative immunofluorescence analysis of TFEB (red, grayscale) subcellular localization in tumor FFPE samples from BHD cohort. Arrowheads indicate tumor cells (nuclear TFEB positive), whereas asterix’ indicate stromal cells (nuclear TFEB negative). Scale bar: 100 µm (main); 25 µm (inserts). (**B**) Quantification of nuclear/cytosolic distribution of TFEB in either stromal (beige) or epithelial tumor cells (blue) from tumor FFPE samples from BHD cohort. (**C**) Representative immunofluorescence analysis of SQSTM1/p62 (magenta) in two separate FFPE samples from BHD cohort (C_i_ = patient case #1-1, C_ii_ = patient case #5). Dotted line indicates border between normal cortex and tumor tissue. Arrowheads indicate tumor cells (nuclear SQSTM1/p62 positive), whereas asterix’ indicate stromal cells (nuclear SQSTM1/p62 negative). Scale bar: 100 µm (main); 20 µm (inserts). (**D**) Quantification of nuclear SQSTM1/p62 spots in tumor and normal kidney FFPE samples from BHD cohort. Error bars are 95% confidence intervals. Specimen 5 contained three distinct tumors and quantifications are annotated accordingly. (**E**) Immunofluorescence analysis of SQSTM1/p62 (magenta) and lysosomal marker LAMP1 (green) in kidney tumor FFPE samples (Sample from BHD patient #4). Representative nuclei outlined in yellow. Scale bar: 20 µm. (**F**) Histogram plot displaying distribution of nuclear SQSTM1/p62 spot diameter (nm) in BHD tumor cells. (**G**) colocalization of SQSTM1/p62 (magenta) and PML (cyan) in kidney tumor FFPE samples (Image shown from BHD patient #4). Nuclei outlined in yellow. Scale bar: 20 µm. (**H-I**) Representative immunofluorescence analysis of SQSTM1/p62 (H, magenta/gray inverted) and TFEB (I, red/gray) subcellular localization in normal kidney cortex FFPE samples from BHD cohort. Scale bar: 100 µm. (**J**) Quantification of nuclear/cytosolic distribution of TFEB in either distal (blue) or proximal tubuli cells (beige) from normal kidney cortex FFPE samples from BHD cohort. Stromal cells and glomeruli were excluded from the analysis. (**K**) Soft agar colony growth for UOK257 cells treated with siRNA targeting *SQSTM1*. Images shown for *SQSTM1* amplicon #2. (**L**) Colony count for soft agar assay for UOK257 cells depleted of SQSTM1/p62 using siRNA targeting *SQSTM1*. Bar plots show mean ± SEM.

**Table 1:** Clinical features of BHD patient samples.

| Patient # | Specimen # | Age range, Sex | Affected kidney | Histological diagnosis and features | Tumor diameter (mm) | Stadium (pTNM) |
| --- | --- | --- | --- | --- | --- | --- |
| BHD1 | BHD1-1 | 55-59, M | Right | Hybrid tumor (with features of chromophobe RCC, oncocytoma and focally of clear cell RCC) | 30 | pT1a |
|  | BHD1-2 (1 year later) | 55-59, M | Left | Multiple (6) hybrid tumors (with features of chromophobe RCC, oncocytoma, and focally of clear cell RCC) | 13; 8; 5; 4.5; 3.5; 1 | pT1a |
| BHD2 |  | 45-49, M | Left | Chromophobe RCC | 20 | pT1a |
| BHD3 |  | 65-69, M | Right | Multiple (3) hybrid tumors (with predominantly clear cell RCC- and focally oncocytoma features) | 25; 5; 3 | pT1a |
| BHD4 |  | 50-54, M | Right | Hybrid tumour (with mainly chromophobe RCC, focally oncocytoma- and clear cell RCC features). | 18 | pT1a |
| BHD5 |  | 45-49, F | Right | T1: Hybrid tumor (with features of oncocytoma and chromophobe RCC)<br>T2: Chromophobe RCC<br>T3: Hybrid tumor (with features of oncocytoma, chromophobe RCC and clear cell RCC) | T1: 130<br>T2: 5<br>T3: 7.5 | T1: pT2b<br>T2: pT1a<br>T3: pT1a |
| BHD6 |  | 65-69, M | Right | Hybrid tumor (with features of chromophobe RCC, oncocytoma and clear cell RCC). | 32 | pT1a |
| BHD7 |  | 55-59, M | Right | Hybrid tumor (with features of clear cell RCC, chromophobe RCC and focally of oncoytoma) | 15 | pT1a |
| BHD8 |  | 60-64, M | Left | Hybrid tumor (with features of oncocytoma, chromophobe RCC, and clear cell RCC) | 65 | pT3a |

Immunofluorescence analysis of LC3 in patient samples displayed overall lower levels in tumor tissue compared to adjacent normal kidney tissue (Fig. S4C-D). Moreover, we found that all BHD patient samples exhibited an accumulation of nuclear SQSTM1/p62 spots in tumor cells, either in the entire tumor or in foci (Fig. 4C-D). As expected, nuclear SQSTM1/p62 foci were found to be negative for the lysosomal marker LAMP1 (Fig. 4E), with diameters ranging from approximately 750 to 1500 µm (Fig. 4F). Together with their nuclear localization, these findings indicate that the SQSTM1/p62 spots were aggregates rather than lysosomal structures. Due to the nature of the spots, we hypothesized that these foci were promyelocytic leukemia nuclear bodies (PML-NBs). PML-NBs are membrane-less, nuclear organelles that are known to be involved in post-translational modifications and nuclear protein sequestration [55]. Notably, SQSTM1/p62 foci colocalized with PML protein (Fig. 4G). These PML and SQSTM1/p62 double-positive structures were exclusively observed in the neoplastic epithelial cells, and not in stromal cells. The latter occasionally contained PML nuclear bodies that were SQSTM1/p62 negative (Fig. 4G, annotated by asterix). Based on this striking nuclear localization in the patient samples, we also investigated nuclear localization of SQSTM1/p62 in the patient-derived UOK257 cells. While all cells showed cytoplasmic SQSTM1/p62 aggregates, nuclear foci of SQSTM1/p62 and PML colocalization was found only rarely in UOK257 cells (Fig. S4F).

The tubular epithelium of the adjacent normal kidney tissue rarely displayed nuclear SQSTM1/p62 puncta compared to tumor cells (Fig. 4D). However, when present, nuclear SQSTM1/p62 puncta was confined to a subset of distal tubular structures (Fig. 4H). Overall, the distal tubuli exhibited lower levels of LC3 staining compared to the remaining tubular epithelium in the normal cortex (Fig. S4D). Interestingly, TFEB nuclear translocation also occurred in distal tubuli whereas it was absent in other tubular structures in cortex (Fig. 4I). To enable compartment-specific quantification, an object classifier was trained to detect proximal and distal tubuli in normal kidney cortex tissue sections (Fig. S4C). The analysis confirmed markedly higher nuclear-to-cytoplasmic ratio of TFEB in distal tubuli structures compared to proximal tubuli (Fig. 4J).

Since SQSTM1/p62 has previously been found to be important for cancer development, we hypothesized that its accumulation in the FLCN-deficient UOK257 cells supports the tumorigenic potential of this cell line. To address this, we silenced SQSTM1/p62 in UOK257 cells, and quantified anchorage-independent growth by soft agar assay, a condition under which UOK257 has been shown to outperform FLCN-restored UOK257 [56]. TFEB depletion, used as a positive control, strongly inhibited the anchorage-independent colony formation of UOK257 (Fig. S4G, I). In contrast, silencing of SQSTM1/p62 did not impair the neoplastic transformation potential of UOK257 cells; instead, it increased both colony count and size (Fig. 4J-K, Fig S4H). Concordant results were obtained for anchorage-dependent growth in standard colony formation assay (Fig S4J-K).

Collectively, BHD tumors exhibited TFEB nuclear translocation and nuclear SQSTM1/p62–PML aggregates. Lastly, SQSTM1/p62 was dispensable for anchorage-independent growth in the UOK257 model, indicating that SQSMT1/p62 accumulation in BHD is symptomatic rather than a driver of tumorigenicity.

## Discussion

In agreement with previous studies on TFEB and/or TFE3 [16,30,32,35,41–43], we found TFEB permanently localized in the nucleus upon loss of FLCN in both cell lines and tumor tissue samples from patients. However, the literature contains conflicting results for MTORC1 activity in relation to FLCN. When FLCN expression is lost or reduced, some have reported increased MTORC1 activity in mice models [21,23,57,58], human BHD tumors [58] and cell lines [22,41,43,59,60], while others observed decreased activity in cell lines [12,16,25,26] or mice [26], and some experiments revealed no or minimal change in cells [32,59] or that differential MTORC1 activity is dependent on cell stress conditions [13]. One study found two categories of MTORC1 activity in renal cysts as measured by pRPS6 staining, where larger cysts showed increased levels, whereas smaller cysts showed decreased levels [24]. Therefore, it has been suggested that the MTORC1 signaling is dependent on context and model. The mechanism behind MTORC1 hyperactivity is proposed to result from the transcriptional upregulation of RAG C/D by TFEB or TFE3 hyperactivation that drives an increase in the canonical MTORC1 targets [21,34]. In our comparisons, the levels of phosphorylated RPS6K, RPS6 and EIF4EBP1 were markedly higher in UOK257. By contrast, the phosphorylation of ULK1 at Ser757 remained unchanged upon FLCN restoration, suggesting that there may be another level of regulation for this site. Consistent with this, prior studies have reported that FLCN loss does not affect ULK1 phosphorylation or kinase activity [37]. We conclude that FLCN loss in UOK257 cells selectively augments anabolic signaling and may not significantly alter MTOR-dependent regulation of the initiation of autophagy.

A few studies have examined the relationship of FLCN to autophagy more directly. In FLCN-deficient HK-2 cells, reduced LC3 flux was detected using the tandem GFP-RFP-LC3 reporter as well as reduced levels of LC3-II [37]. By contrast, our findings from quantitative long-lived protein degradation assays in UOK257 did not reveal changes in bulk autophagy. Moreover, we did not observe differences in LC3 lipidation although an increased fraction of cells with enlarged LC3 puncta was observed. Increased LC3B levels and decreased LC3C levels have been observed both in UOK257 compared to UOK257-2 [38] and in other kidney cancer cell lines depleted for FLCN [25]. In line with this, we observed lower LC3C mRNA levels in UOK257 compared to its FLCN reconstituted counterpart. LC3C seems to exert a tumor suppressing effect in kidney cancer through its involvement in selective autophagy [61,62]. Future studies should define how FLCN loss might regulate selective autophagy processes rather than the bulk autophagy activity assessed here. Collectively, we conclude that bulk autophagy was not altered by FLCN loss in UOK257.

UOK257 cells consistently accumulated SQSTM1/p62, forming large puncta containing SQSTM1/p62 and LC3. In agreement with our results, knockdown of FLCN in HK2 cells causes accumulation of SQSTM1/p62 [37,63]. Furthermore, FLCN deletion in mice kidney loop of Henle cells induces cystogenesis accompanied by strong transcriptional overexpression of SQSTM1/p62 [64]. In a liver-specific context, SQSTM1/p62 protein levels were also increased in a knockout of *Flcn* in mice when fed a methionine/choline deficient diet but not standard chow [65], indicating additional modulation by external cues. SQSTM1/p62, known as sequestome-1 because of its ability to form aggregates [66], binds LC3 family proteins to enable autophagic degradation of selected cargo [51] and can recruit ULK1-complex to induce selective autophagy [67]. SQSTM1/p62 levels are normally controlled by constitutive autophagic degradation, and autophagy impairment can consequently lead to build-up of SQSTM1/p62-containing inclusion bodies [68]. In the case of FLCN-negative BHD cells, our data suggests that the observed SQSTM1/p62 aggregates without an accompanying decrease in autophagy flux was likely due to increased SQSTM1/p62 expression rather than autophagy impairment. Beyond autophagy, SQSTM1/p62 interfaces with proteostatic pathways via its PB1 domain to engage the proteasome and via its UBA domain to clear ubiquitinated proteins and aggregates by aggrephagy [52,69,70]. However, the balance between these functions may be perturbed by a change in overall SQSTM1/p62 levels since overexpression of SQSTM1/p62 can result in large aggregates that mimic autophagic blockade and disrupt the ubiquitin-proteasome system [53]. Whether the observed SQSTM1 accumulation also affects the ubiquitin-proteasome system in BHD would be of interest for future investigations.

Similar to the situation in UOK257, we observed accumulation of SQSTM1/p62 bodies in BHD patient tumor samples. These were also likely not membrane encapsulated due to their nuclear localization and lack of LAMP1 staining. Notably, these aggregates were found in PML nuclear bodies in the tumor cells of the patient samples whereas they were observed predominantly in the cytoplasm in the UOK257 cell line. Although SQSTM1/p62 is generally considered a cytosolic protein, it contains a nuclear localization signal (NLS) and can translocate to the nucleus where it facilitates nuclear protein degradation in PML bodies [71]. It has recently been shown that RCC cell lines overexpress PML, which in turn inactivates p53 and promotes tumor growth [72]. It is therefore possible that SQSTM1/p62 plays a role in tumor development specifically at the PML nuclear bodies, although we could not recapitulate the finding in the UOK257 cell line.

Strikingly, mice with kidney-specific knock-out of FLCN develop renal enlargement, whereas mice with combined kidney-specific knock-out of FLCN and TFEB appeared with no kidney abnormalities, suggesting that TFEB is the main driver behind the development of kidney cancer in BHD syndrome [21]. It is therefore of great interest to understand which of the TFEB-regulated genes are essential for tumor development. As *SQSTM1* is one of the most robustly upregulated gene products upon loss of FLCN in a TFEB-dependent manner [35,64], we hypothesized that SQSTM1/p62 might be essential for tumor development downstream of TFEB. SQSTM1/p62 dysregulation and accumulation are associated with cancerous pathologies including hepatocellular carcinoma [73–75] ovarian [76] and breast cancer [77,78]. Notably, sustained SQSTM1/p62 expression has been shown to promote tumor growth and metastasis in autophagy-deficient cells [79,80], whereas overexpression alone was shown to be insufficient to drive metastatic progression of autophagy-competent breast cancer cells [81]. Consistently, knockdown of SQSTM1/p62 in UOK257 cells, which we found to have functional autophagy, did not affect the tumorigenic potential of this cell line, implying that SQSTM1/p62 may not be pivotal for cancer progression in BHD.

Oncocytoma, chromophobe RCC and hybrid tumors are believed to derive from intercalated and principal cells of the distal tubuli and cortical collecting tubuli [82,83]. Consistent with previously reported findings [35], we also found TFEB in the nucleus in the distal tubuli of the cortex in the adjacent normal kidney tissue of BHD patients. Interestingly, we observed nuclear SQSTM1/p62 and reduced levels of LC3 of distal tubuli cells indicating that these events are co-ocurring. Whether the TFEB and SQSTM1/p62 nuclear positive cells correspond to the intercalated and/or principal cells that eventually give rise to the tumor in BHD should be addressed by future studies.

In conclusion, we found that both cell lines and tumor samples from BHD patients presented with high levels of SQSTM1/p62 in the absence of FLCN. Taken together, our data suggest that FLCN loss primarily alters the balance of TFEB targets rather than affecting bulk autophagy. To which extent dysregulation of other TFEB-regulated genes are essential for tumorigenesis in BHD syndrome warrants further investigation.

## Materials and Methods

### Antibodies

The following antibodies were used for detecting the following targets by immunoblotting and/or immunofluorescence: ACTB/Actin (Abcam, 8227 – 1:1000 WB), FLCN (Cell Signaling Technologies, 3697 – 1:1000 WB), LC3 (MBL, PM036 – 1:500 IF), LC3 (Cell Signaling Technologies, 2775 – 1:1000 WB), phospho-RPS6K Thr389 (Cell Signaling Technologies, 9205 – 1:1000 WB), phospho-TFEB Ser211 (Cell Signaling Technologies, 37681 – 1:1000 WB), phospho-ULK1 Ser757 (Cell Signaling Technologies, 6888 – 1:1000 WB), phospho-EIF4EBP1 Thr37/46 (Cell Signaling Technologies, 2855 – 1:1000 WB), SQSTM1/p62 (MBL, 610833 – 1:16000 WB), SQSTM1/p62 (Progen, P62-C – 1:100 IF), RPS6K (Cell Signaling Technologies, 9202 – 1:1000 WB), TFEB (Abcam, ab122910 - 1:250 IF), TFEB (Cell Signaling Technologies, 4240 – 1:1000 WB), ULK1 (Cell Signaling Technologies, 8054 – 1:500 WB), VCL/Vinculin (Sigma Aldrich, V9131 – 1:2000 WB), EIF4EBP1/4E-BP1 (Cell Signaling Technologies, 9644 – 1:1000 WB).

### Cell culture

Cells were cultured in DMEM/F-12 GlutaMAX (Thermo Fisher Scientific/Gibco 10565018). For UOK257-2 cells 2 µg/ml blasticidin (InvivoGen, ant-bl) was added. All media were supplemented with 10% inactivated FBS (Sigma-Aldrich, F7524) and 1% Penicillin/Streptomycin (Gibco, 15140122). All cells were maintained at 37°C and 5% CO_2_. Cells were routinely checked for mycoplasma contamination and found to be mycoplasma free.

### Cell lysis and western blotting

Cells were seeded overnight in a 6 well plate with 3-4 x 10^6^ cells per well. The cells were washed once with PBS, then lysed in RIPA lysis buffer (Millipore, 20-188) supplemented with phosphatase (Roche, 04906837001) and protease (Roche, 05056489001) inhibitors. The cells were then scraped, transferred to Eppendorf tubes and sonicated using Bioruptor (Diagenode). Thereafter the cells were left for 20 minutes on ice, centrifuged at 14.000 ×g for 10 minutes at 4°C and transferred to new Eppendorf tubes. When preparing lysates that were destined for immunoblotting of phosphorylated proteins, the protocol was shortened by omitting the sonication and incubation steps to preserve the phosphorylated sites. The protein concentration was measured with Pierce Bicinchoninic Acid Assay (BCA) Protein Assay Kit (Thermo Fisher Scientific, 23227).

Proteins were mixed with 4X Nu-Polyacrylamide gel electrophoresis (NuPAGE) LDS Sample Buffer 4x (Invitrogen, NP0008) and 10X 1M DL-Dithiothreitol (DTT) (Sigma-Aldrich, D0632) to a final concentration of 1X, and boiled at 95°C for 10 minutes. The samples were loaded on a 4–20% Criterion™ TGX™ Precast Gel (Bio-Rad, 5671093/5671094/5671095) and separation of protein by SDS gel-electrophoresis with 80V-150V. The proteins were transferred to a 0.45 µm LF PVDF membrane (Bio-Rad, 161–0374) using semi-dry Trans-Blot® Turbo™ Transfer System (Bio-Rad) for 7 minutes at 25V. Membranes were blocked for 10 minutes by air drying. Membranes were then incubated with primary antibodies in TBS with 0.1% Tween-20 (Sigma-Aldrich, P1379) (TBS-T) and 5% Bovine serum albumin (BSA) (Sigma-Aldrich, A2153) overnight at 4°C or 2 hours at room temperature. Afterwards, the membranes were washed three times for 10 min in TBS-T and then incubated for 1 hour at room temperature with secondary antibodies. The secondary antibodies that were used was: HRP-donkey-anti-goat, 1:5000 (Jackson, 705-035-147), HRP-goat-anti-rabbit, 1:5000 (Jackson, 111-035-144), HRP-goat-anti-mouse, 1:500 (Jackson, 115-035-003).

The immunoblots were visualized using SuperSignal west pico/dura/femto Chemiluminescent Substrate (Thermo Fisher Scientific, 34580/34076/34095) with either Chemidoc (Bio-Rad, Hercules, CA, USA) or Sapphire™ Biomolecular Imager (Azure biosystems). Quantification was performed using ImageLab 6.1 Software (Bio-Rad). The intensity of each band was measured and the background intensity was subtracted using the Global Subtraction Method.

### Drug treatment and starvation

Cells were starved for 3 hours in EBSS (Thermo Fisher Scientific, 24010043) unless specifically stated otherwise. Cells were treated with drugs for 3 hours unless specifically stated otherwise. The drugs and concentrations used were DMSO (Sigma-Aldrich, D2650), SAR405 (Selleckchem, S7682) 5 µM (WB, IF, flow cytometry); 10 µM (LLPD), BafA1 (Enzo Life Sciences, BML-CM110-0100) 100 nM (WB, IF, LLPD), Chloroquine (Fluka BioChemika, 25745) 100 µM (flow cytometry), Torin 1 (Tocrin Bioscience, 4247) 250 nM (IF, WB, flow cytometry), AUTEN99 (Cayman Chemical, 30047) 10 µM (WB, flow cytometry), MRT68921 (Sigma-Aldrich, SML1644) 4 µM (WB, flow cytometry).

### Electron microscopy

For immunolabelling and EM cells were prepared as described earlier [84,85]. In brief, cells were fixed with 4% PFA (Polysciences, 18814-20) and 0.1 M glutaraldehyde in 0.1 M PHEM buffer (60 mM PIPES, 25 mM HEPES, 2 mM MgCl_2_, 10 mM EGTA; pH 6.9) overnight. The cells were embedded in gelatin, infused with 2.3 M sucrose, frozen and cryosectioned before subjected to immunolabeling with rabbit anti-SQSTM1/p62 (MBL, PM045) and protein A conjugated 10 nm gold particles. Images were obtained with 80kV in JEOL-JEM 1230 electron microscope equipped with Morada camera (Olympus, Germany) and iTEM software.

### Immunofluorescence and confocal imaging of cells

Cells were seeded overnight in an 8 well plate (Ibidi, 80827) with 60 000 cells per well. The cells were washed once with PBS, then fixed with 4% paraformaldehyde (Polysciences, 18814-20) for 15 minutes at room temperature. Then they were washed with PBS, quenched for 10 minutes using PBS with 0.05 M NH_4_Cl, permeabilized for 5 minutes using PBS with 0,5% Triton TX-100 (Sigma-Aldrich, 9002-93-1). Then the cells were stained with primary antibodies for 1 hour at room temperature. Thereafter the samples were washed 3 times for 5 minutes with PBS with 0,5% saponin before incubation with secondary antibodies for 1 hour at room temperature. The secondary antibodies that were used were donkey-anti goat A555 1:500 (Molecular Probes, A21432), donkey anti-rabbit A488 1:200 (Jackson, 711-545-152), donkey anti-guinea pig Cy3 1:500 (Jackson, 706-165-148), donkey anti-mouse A647 1:500 (Jackson, 715-605-151). Thereafter the samples were washed once for 5 minutes with PBS with 0,5% saponin, followed by incubation with 1 µg/mL Hoechst 33342 (Thermo Scientific, 62249) for 10 minutes, 5 minutes of washing with PBS with 0,5% saponin and one final wash with PBS. The cells were left in PBS or in ProLong Diamond Antifade Mountant (Invitrogen, P36961) at 4°C. Imaging was performed with Nikon Yokogawa CSU-W1 40x air (NA 0.95, Air) and laser set of 405 nm, 488 nm and 561 nm, with a z-stack covering 11/15 µm with a z-step of 0,6 µm.

### Image processing and quantification

Within each set of experiments, images were captured with identical settings below pixel value saturation and post-processed identically. Intensity enhancements for visualization purposes on confocal microscopy images were performed using Adobe Photoshop, ImageJ or NIS Elements, and all intensity alterations were done equally for all samples to ensure comparativeness within each experiment unless otherwise stated. Confocal images shown in figures are maximum intensity projections made using the NIS Elements software unless otherwise stated. Image quantifications for confocal images were either performed with NIS Elements software or Fiji/ImageJ. Image processing and quantifications for patient samples were performed with QuPath [54] and Fiji/ImageJ.

### Birt-Hogg-Dubé patient samples

BHD patients who had previously undergone surgery for kidney tumor were identified by using the clinical database at the Department of Medical Genetics, Oslo University Hospital. Surgical specimens with kidney tumor of eight non-related BHD patients were collected from different hospitals in the Health Region South-Eastern Norway after written informed consent was obtain from all of them. The use of these samples in this study has been approved by the Regional Committee for Medical Research Ethics (REK South-East Norway, #48926) and by the Data Protection Officer at Oslo University Hospital (Reference number 20/22703). All surgical specimens were reviewed by experienced histopathologist and cases were classified according to the WHO-2022 classification. The age and gender of the patients, side of the lesion, histological types, morphological composition, tumor size and pTNM stage are summarized in Table 1. Morphological and immunohistochemical features of the tumors are shown in Table S1.

### Immunofluorescence of patient samples

Serial sections of FFPE tissue samples were cut at 5 µm thickness and placed on SuperFrost™ Plus glass slides (Epredia). To perform immunofluorescence staining, FFPE slides were heated at 60°C for 10 minutes before deparaffinization by submerging the slides in the following solutions: Xylene (Sigma Aldrich, 534056) 2 x 5 minutes, 99,5% EtOH for 2 x 3 minutes, 95% EtOH for 1 minute and 70% EtOH for 1 minute. Slides were boiled in with 10 mM Tris-EDTA buffer (pH 9,4) in a rice cooker for 30 minutes for antigen retrieval. Slides were washed in PBS with 0,1% Tween (PBS-T) twice for 3 minutes. Slides were blocked with blocking solution of Donkey serum 5% (Abcam, ab7475 Lot# GR3389768-1) in 0,1% PBS-T for 30 minutes, before they were stained with primary antibodies diluted in blocking solution overnight at 4 degrees with slight tilting. Primary antibodies and dilutions in this study were as follows: TFEB (Cell Signaling Technologies, 37785) 1:300, SQSTM1/p62 (Abnova, H00008878-M01 batch K8251-2C11) 1:300, 1:100, LAMP1 (Sigma, L1418, batch 0000135033) 1:300, PML (Cell Signaling Technologies, 69789S, batch 1) 1:300 (FFPE tissue)/1:500 (cells). After incubation with primary antibodies, slides were washed with PBS-T for 5 minutes 3 times. Slides were then incubated with secondary antibodies diluted in PBS-T for 3 hours at room temperature in the dark. Secondary antibodies and dilutions were as follows: donkey anti-rabbit Alexa-568 (Molecular Probes, A10042, LOT 1891789) 1:500, donkey anti-mouse Alexa-568 (Molecular Probes, A10037) 1:500, donkey anti-rabbit Alexa-647 (Jackson ImmunoResearch, 711-605-152, LOT 159933) 1:500 and donkey anti-mouse Alexa-647 (Jackson ImmunoResearch, 715-605-150, LOT 151088) 1:500. Slides were washed twice with PBS-T, stained for nuclei with Hoechst (Thermo Scientific, 62249) for 10 minutes and mounted with 1 drop of ProLong Diamond Mounting Media (Invitrogen, P36961) per tissue section. The immunostained sections were imaged on either OLYMPUS VS200 Slide Scanner equipped with an X-cite XYLIS LED Fluorescence lamp (Excelitas Technologies®) or Nikon ECLIPSE Ti2 microscope equipped with CrestOptics X-light V3 confocal spinning disk. HE/HPS sections were imaged with OLYMPUS X-line Apo 20X/NA0.8 objective and immunofluorescence samples with OLYMPUS X-line Apo 40X/NA0.95 objective, Nikon Plan Apo λD 40X/NA0.95 air objective or Nikon Plan Apo λD 60X oil objective. Immunofluorescence images were captured with a Z-stack of 4-6 µm. HE/HPS images were captured with a single Z-layer. Immunostained sections were annotated by a combination of HE/HPS staining and Hoechst staining. 4000-8000 cells in representative areas of both normal and tumor tissue were evaluated for each case. BHD patient case #7 and #8 were omitted from immunofluorescence analysis due to the quality of the tissue preservation being insufficient for those analyses. In tumor tissue, stromal cells, connective tissue and cystic areas were excluded from the analysis. For normal tissue, only tubule cells from the cortex of the kidney were included in analysis. Medulla, glomeruli and stromal cells were excluded. Areas damaged by diathermy and normal areas adjacent to tumor, up to a distance of 0.5 mm, were excluded in the analysis. QuPath was used to segment cells and measure signal intensities [54]. Detection of epithelial tumor- and tubule cells in tumor and normal tissue respectively was automated with a trained object classifier in QuPath, which was validated by a trained pathologist. SQSTM1/p62 nuclear spot count was performed in Fiji/ImageJ after cell segmentation and classification in QuPath.

### Long lived protein degradation assay (LLPD)

We used a modified version of the method described by Luhr and colleagues [50]. The cells were seeded and incubated in 0,135 µCi/ml L-(^14^C)-valine (Perkin Elmer, NEC291E) supplemented RPMI 1640 complete medium (Thermo Fisher Scientific/ Gibco, C21875091) for 24 hours. Proceeding, a 3-hour chase in RPMI complete medium supplemented with 10 mM L-Valine (Sigma-Aldrich, V0500), which is not radioactive, to let short lived proteins degrade. The medium was removed and the cells were then incubated in complete medium or EBSS with 10 mM valine with either 100 nM BafA1 or 10 µm SAR405 for 4 hours. Thereafter, we used 50% trichloroacetic acid (TCA) (Merck, 1.00807.1000) for the precipitation of cellular proteins, and it was incubated overnight. 0.2 M KOH (Merck, 221473) was added to solubilize the solution and the percent degradation was calculated as the acid-soluble radioactivity divided by total radioactivity. To induce autophagy, we used EBSS starvation (Thermo Fisher Scientific, 24010043), and to calculate the autophagic contribution to the degradation of the long lived protein we used two autophagic inhibitors: SAR405 (Selleckchem, S7682) which prevents autophagosome formation by inhibiting PIK3C3, and BafA1 (Enzo Life Sciences, BML-CM110-0100) which prevents fusion of the autophagosome and the lysosome by inhibiting the v-ATPase.

### Nuclear/cytosolic fractionation

We used a modified version of the method described by Senichkin and colleagues [86]. The cells were washed with ice cold PBS, then lysed with lysis buffer (50 mM Tris-Cl (pH7,5), 150 mM NaCl, 5 mM EGTA, 10% glycerol, 0,1% NP-40), incubated for 10 minutes on ice and centrifuged at 1000 x g at 4°C for 5 minutes. The supernatant and pellet were each transferred to a new Eppendorf. The pellet (the nuclear fraction) was resuspended in the isotonic buffer and incubated on ice for 10 minutes. The Eppendorfs were then centrifuged at 1000 x g at 4°C for 3 minutes. The supernatant was removed, and this washing was repeated three times. Finally, the nuclei were resuspended in 50 µl lysis buffer and sonicated using the Bioruptor (Diagenode). The supernatant from the first step (the cytoplasmic fraction) was centrifuged at 15000 x g at 4°C for 3 minutes, and the supernatant was transferred to another Eppendorf. Samples were then subjected to western blotting as described above.

### Phospho flow cytometry

Phospho flow cytometry protocol was adapted from previous reports [87]. Cells were seeded overnight in a 6 well plate with 300 000 cells per well. The next day drugs were added for 3 hours. Cells were brought into single cell suspension with Accutase (Sigma-Aldrich, A6964) treatment for 45 min to avoid loss of phosphorylation. Cells were centrifuged at 500 x g at 4°C for 5 minutes, and the pellet was washed once with PBS and resuspended in 500 µl PBS. Cells were fixed by adding 16% PFA (Polysciences, 18814-20) to a final concentration of 1,5% PFA at 37°C for 10 minutes. Fixed cells were washed twice with PBS, before 200 µl of ice-cold BD Phosflow Perm Buffer III (BD Biosciences, 558050) was added to each sample drop-wise while vortexing to avoid cell clumping. Cells were directly transferred to -80°C for a minimum of 30 minutes. Permeabilization occurred prior to fluorescent cell barcoding (FCB) to enable proper penetration of the fluorescent NHS-dyes to improve separation of cell populations. Ice-cold PBS was added to cell suspensions and cells were added to 96-well V-bottom plate (Thermo Scientific, 249662, Lot#163248) containing 30 µl of FCB dye combination of Alexa 488 succinimidyl ester (Invitrogen, A20000, Lot#2272578), Pacific Blue succinimidyl ester (Invitrogen, P10163, Lot#2433885) and Pacific Orange succinimidyl ester (Invitrogen, P30253, Lot#2025923) diluted in DMSO and stained for 20 minutes in the dark at room temperature. Details on FCB dye concentrations, combinations and controls can be found in Table S2. Cells were washed three times with flow wash (PBS + 5% FBS) and resuspended in 250 µl flow wash. Barcoded cell populations were combined in a 15 ml tube, centrifuged at 500 x g at 4°C for 5 minutes and resuspended in the amount of flow wash which allowed for 25 µl of cell suspension per phospho-antibody. Single FCB dye compensation controls were kept in separate tubes. 25 µl of combined cell suspension were added to 0,5 ml Eppendorf tube containing 25 µl Alexa-647-conjugated phospho-antibody. An overview of antibodies can be found in Table S3. Cells were stained for 30 minutes in the dark at room temperature. Samples were washed twice with flow wash, resuspended in 400 µl flow wash and filtered in a FACS tube with 35 µm filter cap (Falcon, 352235). Samples were run on a LSRII UV laser flow cytometer with FACSDiva software. Single FCB dye compensation controls were used to calculate the compensation matrix prior to each run. The background signal for UOK257-1 and UOK257-2 was measured as equal. Cell populations were deconvoluted based on FCB signal and single cell population signal of Alexa 647 was analyzed with FlowJo and reported as MFI (Median Fluorescent Intensity) after isotype normalization and global scaling to adjust for technical variations between experiments. Results were analyzed with R/R Studio.

### siRNA transfection

Lyophilized siRNA was reconstituted with nuclease-free water and stored at -20°C. For reverse transfection, transfection complexes were prepared by combining DMEM free from serum and antibiotics (Gibco, 61965 Lot# 2533742), siRNA, and Lipofectamine RNAiMax (Invitrogen, 13778030, Lot# 2946891) according to manufacturer’s instructions and incubated at room temperature for 15-20 minutes. siRNAs used in this study were siRNA Silencer Select Negative Control No. 1 (Thermo Fisher Scientific, 4390844), *TFEB* siRNA s15496 (#1 in text) and s15497 (#2 in text) (Life Technologies, 4392420), *SQSTM1* siRNA s16960 (#1 in text) and s16962 (#2 in text) (Ambion). Cell suspensions of UOK257-1 and UOK257-2 were prepared in their respective complete media and mixed with corresponding siRNA transfection complexes to a final concentration of 20 nM siRNA. The mixtures were seeded into appropriate wells and incubated at 37°C with 5% CO_2_ for 48 hours.

### Soft agar colony formation assay

For the soft agar colony formation assay, 5% agar solution was prepared by mixing 5 g agar (Sigma, A1296-100G, Lot# 8C8R1697V) with 100 mL MQ water and autoclaved at 121°C for 15 minutes. The sterile solution was maintained at 50°C during experiments to preserve liquid phase. The bottom agar layer was prepared by adding 750 µl of 0,7% agar in complete media to wells of 12-well plates, which was allowed to solidify at 4°C for at least 15 minutes followed by drying in a clean hood for 30 minutes without lid to avoid separation of agar layers. siRNA transfected cells were brought into suspension by trypsinization, and 750 µl of 0,4% agar solution containing 1,500 cells were added on top of the solidified bottom gel layer and allowed to solidify for 30 minutes in room temperature. Plates were incubated at 37°C with 5% CO₂, with 750 µl complete medium added on top as feeding layer the following day. Feeder layer was refreshed weekly. After 4-6 weeks, medium was removed, wells were washed with 500 μl PBS, and colonies were stained with 400 µl of 1 mg/mL MTT solution (Invitrogen, M6494 Lot# 2936400) at 37°C with 5% CO_2_ for 4 hours before imaged and automatically counted using the “Find Maxima” command in Fiji/ImageJ.

### Primers

Primers used in this study are listed in the table below. Efficiency of primer sets was tested before running expression level analysis.

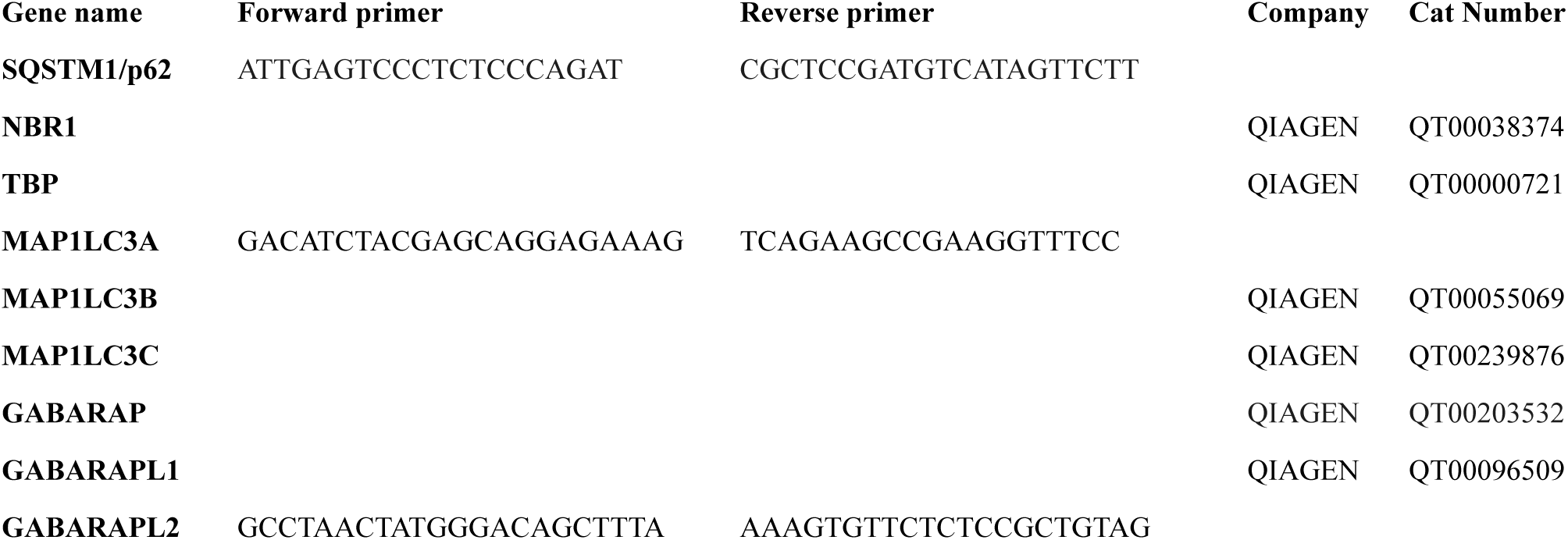

### RNA extraction, reverse transcription and quantitative PCR

Transcriptional levels of genes of interest were assessed by RT-qPCR. For this, UOK257 and UOK257-2 cells were washed once in PBS1X and frozen at -80°C. QIAGEN RNeasy Plus lysis buffer was added to collected cells and RNA extraction was performed using RNA extraction kit (QIAGEN RNeasy Plus, 74134). Upon isolation, RNA concentration was measured by NanoDrop 1000 Spectrophotometer. Complementary DNA (cDNA) was synthesized from RNA through reverse transcription, accordingly to the iScript™ cDNA Synthesis Kit (BIO-RAD, 1708891) using the thermocycler (Applied Biosystems SimpliAmp PCR machine, A24811). RT-qPCR was carried out with Fast SYBRGreen Master Mix (Applied Biosystems, 4385614) on the Applied Biosystems StepOnePlus Real Time PCR system (Applied Biosystems, Carlsbad, CA, USA), with StepOne v2.3 software. The 2^−ΔΔCt^ method was used to measure the relative expression levels.

### Statistical analysis/ Experimental design and statistics

Statistical analysis is indicated in the legends for each figure panel.

### AI Usage Statement

The preparation of this scientific manuscript involved the use of an AI language model, specifically OpenAI’s GPT-4, to assist in drafting, revising, and enhancing the clarity of the text. The model did not perform data analysis, generate results, or interpret findings. No patient data was fed to the model. All content generated by the model was thoroughly reviewed and verified by the authors, who retain full responsibility for the accuracy and integrity of the text.

## Acknowledgments

We thank W. Marston Linehan, NCI Bethesda, for generously sharing the UOK257 and UOK257-2 cell lines. We thank S. Skånland for providing antibodies and assistance with the phospho-flow protocol. The Oslo University Hospital core facility for Advanced Electron Microscopy Montebello node, the core facility for Advanced Light Microscopy Montebello node, the core facility for flow cytometry Montebello node, and the MolMed Imaging Platform at University of Oslo are acknowledged for access, help, and services.

HU and FK are part of The Medical Student Research Program (MSR) at the Faculty of Medicine, University of Oslo, and were supported by grant 271555/F20 from the Research Council of Norway and running costs from the Faculty of Medicine. H.K. was supported by grant 30078 from the Research Council of Norway, grants 2017062 and 2022019 from the South-Eastern Norway Regional Health Authority and by the European Union (ERC, FINALphagy, 101039174). FINALphagy is a project in the Centre for Digital Life Norway, which is supported by the Research Council of Norway’s grant 320911. J.M.E. was supported by South-Eastern Norway Regional Health Authority (HSØ grants 2017064, 2018012, 2019049 and 2019096), the Norwegian Cancer Society (grants 182524 and 208012) and the Research Council of Norway through research grants 261936, 294916 and 314811, and from the European Union’s Horizon 2020 research and innovation programme under grant agreement No. 847912 (RESCUER). The research leading to these results has received funding from the European Union’s Horizon 2020 research and innovation programme under the Marie Skłodowska-Curie grant agreement No 801133. This work was partly supported by the Research Council of Norway through its Centres of Excellence funding scheme, project number 262652.

## Author contributions

Halvor Ullern: Formal analysis, Investigation, Writing—original draft, Writing—review & editing, Visualization.

Julie Aarmo Johannessen: Formal analysis, Investigation, Writing—original draft, Writing—review & editing, Visualization.

Feyza Kasikci: Investigation, Writing—review & editing

Miriam Formica: Investigation, Writing—review & editing

Naghmeh Karimi Melve: Investigation, Writing—review & editing

Siri Andresen: Investigation, Writing—review & editing

Andreas Brech: Investigation, Writing—review & editing, Visualization.

Karol Axcrona: Resources, Writing—review & editing,

Kjersti Jørgensen: Resources, Writing—review & editing,

Lorant Farkas: Formal analysis, Investigation, Resources, Writing—review & editing, Visualization.

Jorrit M. Enserink: Writing—review & editing, Supervision.

Helene Knævelsrud: Conceptualization, Formal analysis, Investigation, Writing—original draft, Writing—review & editing, Supervision, Project administration, Funding acquisition.

## Abbreviations

BafA1: Bafilomycin A1
BHD: Birt-Hogg-Dubé
EM: electron microscopy
FLCN: folliculin
FNIP: folliculin-interacting protein
GABARAP: gamma-aminobutyric acid type A receptor-associated proteins
GAP: GTPase Activating protein
IF: immunofluorescence
LC3: microtubule-associated proteins 1A/1B light chain 3
LLPD: Long lived protein degradation
MTOR: mechanistic Target of Rapamycin
PE: phosphatidylethanolamine
PML: Promyelocytic leukemia
PML-NBs: PML nuclear bodies
RAG: Ras related GTP-binding protein
RCC: renal cell carcinoma
RHEB: Ras homolog enriched in brain
TFE3: Transcription Factor E3
TFEB: Transcription Factor EB
WB: western blot

