## Supplemental figures and tables for "SQSTM1/p62 accumulation is a hallmark of FLCN loss in Birt-Hogg-Dubé syndrome-associated kidney cancer"

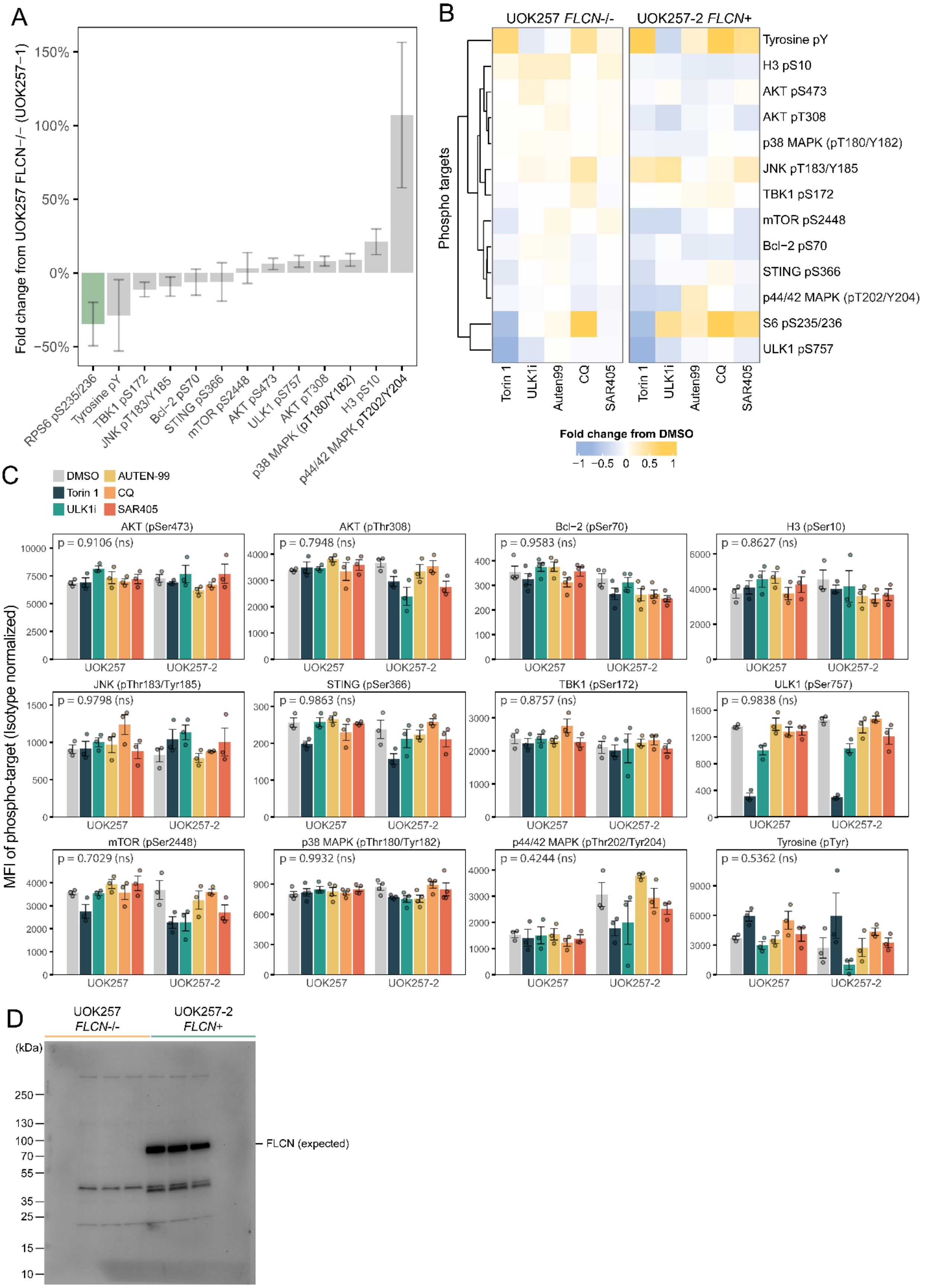


**Figure S1.** Phospho flow cytometry of UOK257 cell lines. (**A**) Waterfall plot showing fold change (%) of phosphorylation levels from UOK257 *FLCN*-/- DMSO. (**B**) Heatmap from phospho flow cytometry, showing log fold change from DMSO for each cell line**.** Yellow indicates increased, while blue indicates decreased phosphorylation levels compared to DMSO. Clustering of phospho targets was performed by Euclidean distance calculation followed by complete hierarchical clustering. (**C**) Bar plots showing normalized median fluorescent intensity (MFI) values of phospho flow cytometry (N=3-4 replicates). P-value on plot indicate results of one-way ANOVA test of overall treatment-specific response in-between cell lines. (**D**) Complete membrane for FLCN immunoblotting. ns: not significant ($P > 0.05$); $*P < 0.05$, $**P < 0.01$, $***P < 0.001$, $****P < 0.0001$.

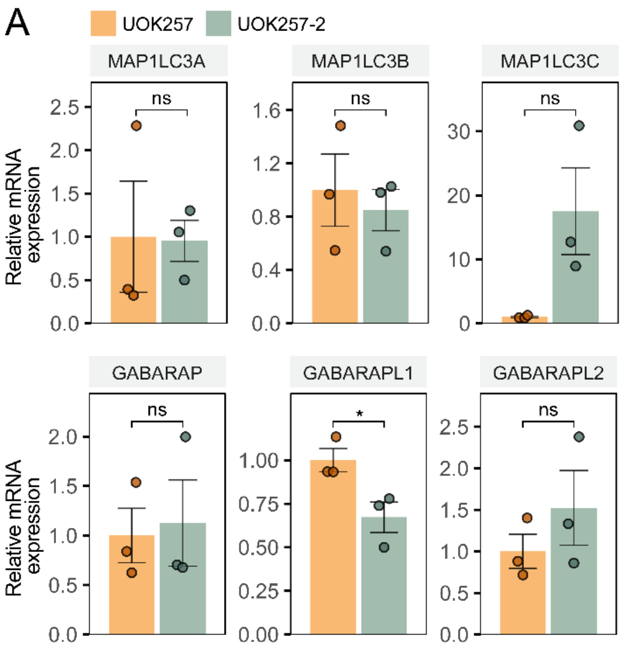


**Figure S2.** Gene expression of some LC3 protein family members is altered by FLCN reconstitution. (**A**) Relative mRNA expression levels of LC3 protein family members measured by RT-qPCR in UOK257 cells. Bar plots show mean ± SEM (N=3). Statistical analysis was performed using two-sided t-test. ns: not significant ($P > 0.05$); *$P < 0.05$, **$P < 0.01$, ***$P < 0.001$, ****$P < 0.0001$.

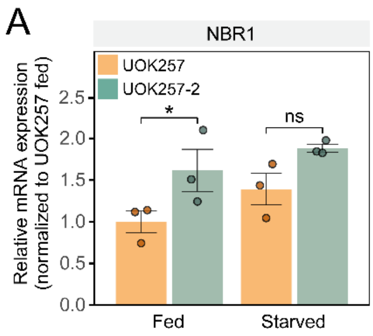


**Figure S3:** NBR1 mRNA expression is decreased in UOK257 cells deficient in FLCN. (**A**) Relative mRNA expression levels of NBR1 measured by RT-qPCR in UOK257 cells. Expression levels are normalized to UOK257 *FLCN-/-* fed sample. Statistical analysis was performed using one-way ANOVA with Tukey’s multiple comparisons test.

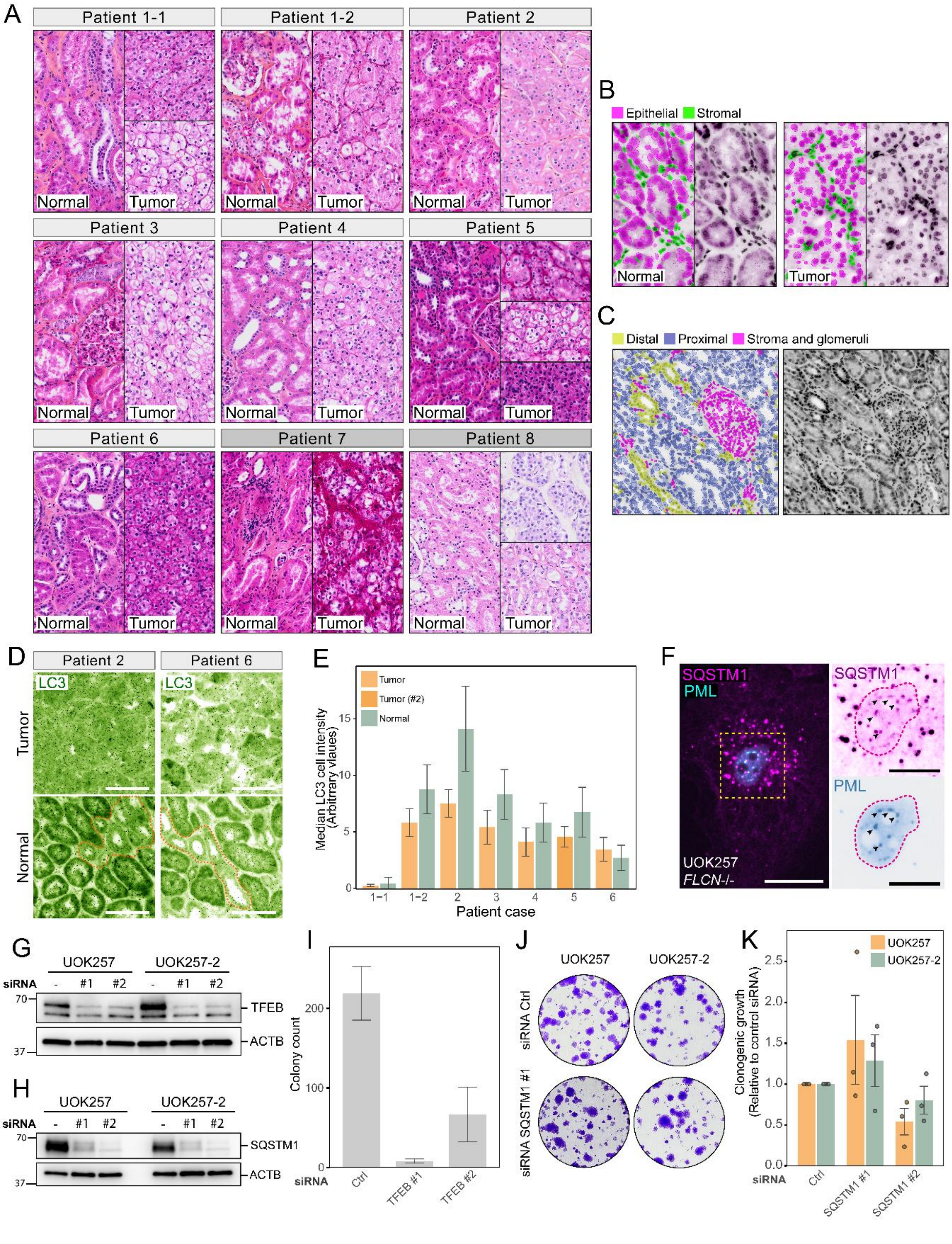


**Figure S4.** (**A**) Representative HE or HSE staining of kidney tumor and normal kidney FFPE samples in BHD cohort. Where multiple tumors were present or the primary tumor presented multiple distinct morphologies, several tumor images are included to illustrate the heterogeneity. Patient samples 7 and 8 (dark grey headers) were omitted from analysis due to insufficient tissue section quality. (**B**) Illustration of trained object classifier in QuPath used to distinguish epithelial (magenta) and stromal (green) cells in normal and tumor tissue for quantification purposes. Nuclei in black, TFEB in purple. Note that TFEB signal is just present in figure to visualize the structures, the classifier was trained on nuclei features only. (**C**) Illustration of trained object classifier in QuPath to distinguish distal tubuli (yellow), proximal tubuli (blue) and stroma/glomeruli (magenta) in normal kidney cortex tissue samples. (**D**) Immunofluorescence images of LC3 in kidney tumor and normal kidney FFPE samples in BHD cohort. Distal structures are outlined in orange. Scalebar: 100 µm. (**E**) Bar plot showing median LC3 intensity in epithelial tumor or normal kidney cells as shown in panel E. For patient case #5, only tumor sample #2 was analyzed. (**F**) Immunofluorescence images of UOK257-1 cells showing nuclear colocalization of SQSTM1/p62 (magenta) and PML (cyan). Black arrowheads indicate nuclear colocalization puncta. Scale bar: 20 µm (main panel, white) and 10 µm (inserts, black). (**G-H**) Immunoblot demonstrating efficiency of siRNA targeting *TFEB* (G) and *SQSTM1* (H) in UOK257 cells. 2 amplicons were used per target. (**I**) Soft agar assay for UOK257 *FLCN*-/- cells with *TFEB* siRNA treatment. (**J**) Representative images of colony formation assay for UOK257 and UOK257-2 treated with siRNA targeting *SQSTM1*. (**K**) Quantification of clonogenic growth from colony formation assay. Clonogenic growth was quantified as percentage (%) of growth area covered and normalized to siRNA Control respectively for each cell line.

**Table S1:** Extended histological features of BHD specimens

| Patient # | Specimen # | Histological diagnosis and features | Morphological features of tumor | Tumor immunophenotype |
|---|---|---|---|---|
| BHD1 | BHD1-1 | Hybrid tumor (with features of chromophobe RCC, oncocytoma and focally of clear cell RCC) | **Growth pattern:** Mainly solid, in places nested. Focally thin bands of vascularized connective tissue.<br>**Cell morphology:**<br>Cytoplasm: Mainly pale reticular eosinophilic cytoplasm. Patchy areas with oncocytic, eosinophilic cytoplasm, while in other areas clear cytoplasm. Occasionally perinuclear halo is present. Cell borders vary between invisible and weakly/ moderately marked. In the clear cell areas, the cell borders are prominent. Colloidal blue stain is weakly positive in the cytoplasm in some places.<br>Nucleus: Mainly round nuclei with inconspicuous, occasionally moderately marked nucleoli. Occasional binucleated forms are present. Raisinoid nuclei are not a feature. | CD117+, CK7-/focally+, CK20-, EMA-/+, CD10+/focally-, vimentin-, CA IX-, AMCR-, s100- |
| | BHD1-2 | Multiple (6) hybrid tumors (with features of chromophobe RCC, oncocytoma, and focally of clear cell RCC) | **Growth pattern:** Mainly solid, in places nested, very focally vaguely papillary pattern. Often collagenous bands of connective tissue.<br>**Cell morphology:**<br>Cytoplasm: Mainly pale reticular eosinophilic cytoplasm. Focally oncocytic eosinophilic cytoplasm, while in other small areas clear cytoplasm. Perinuclear halo is present in places. Cell borders are mainly weakly to moderately marked, while in other areas they are invisible; and in the clear cell areas they are prominent. Colloidal blue stain is weak positive in the cytoplasm in places.<br>Nucleus: Mainly round nuclei with mainly inconspicuous, occasionally moderately marked nucleoli. Occasional binucleated forms are present. Raisinoid nuclei are not a feature. | CD117+, CK7-/focally+, CK20-, EMA+, CD10-/+, vimentin-, CA IX-, AMCR+/-, S100-. |
| BHD2 | | Chromophobe RCC | **Growth pattern:** Mainly solid, focally cystic, papillary or tubular patterns. Thin, hyalinised septa in some areas. In some areas fresh hemorrhage and also hemosiderin macrophages.<br>**Cell morphology:**<br>Cytoplasm: Mainly pale-reticular to eosinophilic cytoplasm. Clear perinuclear halo throughout the tumor. Cell borders are from moderately to prominently marked.<br>Nucleus: Mainly raisinoid/ focally round nuclei with occasional inconspicuous nucleoli. Frequent binucleated forms are present. | CD117-, CK7-/+, PAX8+, CD10-, vimentin-, CA IX- |
| BHD3 | | Multiple (3) hybrid tumors (with predominantly clear cell RCC- and focally oncocytoma features) | **Growth pattern:** Predominantly solid, focally nested/ rosette growth in a myxoid stroma.<br>**Cell morphology:**<br>Cytoplasm: Mainly clear or pale reticular cytoplasm with perinuclear halo. In small areas eosinophilic oncocytic cytoplasm. Cell borders are from moderately to prominently marked except the oncocytic areas where borders are mainly weakly marked/ invisible.<br>Nucleus: Mainly round nuclei with occasional tiny nucleolus. Frequent bi- or multinucleated forms. Focal pleomorphism is noted. | Not performed |

| | | | | |
|---|---|---|---|---|
| BHD4 | | Hybrid tumour (with mainly chromophobe RCC, focally oncocytoma- and clear cell RCC features). | **Growth pattern:** Mainly solid, focally cystic, focally papillary patterns. Focally septa with vascularized connective tissue. Focal hemorrhagy.<br>**Cell morphology:**<br>Cytoplasm: Mainly pale reticular with perinuclear halo and in some areas eosinophilic oncocytic cytoplasm and focally clear cytoplasm. Cell borders are moderately marked except in the oncocytic areas where cell borders are weakly marked/ invisible.<br>Nuclei: Mainly round, focally irregular/ raisinoid nuclei; often with central nucleoli that are inconspicuous to moderately marked. | CD117+; CK7-/+; PAX8+; CD10-/occasionally weakly+; AMACR+/-; vimentin-; CA IX-; melanA-; HMB45-; calreticulin-; Ki67 <1% |
| BHD5 | | T1: Hybrid tumor (with features of oncocytoma and chromophobe RCC)<br>T2: Chromophobe RCC<br>T3: Hybrid tumor (with features of oncocytoma, chromophobe RCC and clear cell RCC) | T1: **Growth pattern:** Mainly solid, focally tubular and trabecular. Focal areas with haemorrhage. In the tubular/ trabecular areas there is acellular pale extracellular matrix. Foamy macrophages are focally present.<br>**Cell morphology:**<br>Cytoplasm: Mainly eosinophilic, in places pale reticular with clear perinuclear halo. Cell borders mainly invisible.<br>Nucleus: Round to slightly irregular contour; small to medium in size with clear chromatin. Very often central nucleolus that is small to moderately marked. Bi- and multinucleated forms are frequent.<br>T2: **Growth pattern**: Tubular and cystic.<br>**Cell morphology:**<br>Cytoplasm: Pale reticular, in places eosinophilic. Prominent perinuclear halo. Cell borders are moderately marked.<br>Nucleus: Mainly round, occasionally slightly irregular. Clear chromatin. Inconspicuous nucleolus.<br>T3: **Growth pattern:** Solid.<br>**Cell morphology:**<br>Cytoplasm: Mainly pale reticular to eosinophilic; clear in some areas. Cell borders vary between invisible and moderately marked. Perinuclear halo is present focally.<br>Nucleus: Mainly round, often with central nucleolus that is small to moderately marked. Raisinoid nuclei are not a feature Occasional bi- and multinucleated forms. | T1: CD117-/f+; panCK(AE1/AE3)+; CK7-/f+; CK8/18+; CK20-; AMACR-; CA IX-; CD10-; vimentin-; E-cadherin-/+; TFE3-; PAX8+; ALK-; cathepsinK-; DOG1-; melanA-; HMB45-; maintained SDHB and FH.<br>T2: CD117+; panCK(AE1/AE3)+; CK7-/+; CK8/18+; CK20-; AMACR+; CA IX-; CD10-/f+; vimentin-; E-cadherin+; TFE3-; PAX8+; ALK-; cathepsinK-; DOG1-; MelanA-; HMB45-; maintained SDHB and FH.<br>T3: CD117+/-; CK7-/+; vimentin-; CA IX-; CD10-/f+ |
| BHD6 | | Hybrid tumor (with features of chromophobe RCC, oncocytoma and clear cell RCC). | **Growth pattern:** Mainly solid and in places trabecular. Focally nested growth in a loose acellular matrix. Focally also papillary pattern.<br>**Cell morphology:**<br>Cytoplasm: Mainly pale reticular, in places oncocytic eosinophilic and focally clear. Cell borders varying between invisible, moderately marked and prominent.<br>Nucleus: Mainly small to medium, in places large. Mainly round nuclei with clear chromatin, but focally raisinoid. Often central nuclei that are from small to moderately marked. Bi- and multinucleated cell forms are present. | CD117+; CK7-/f+; panCK +(dot-like)/+ (entire cytoplasm); CD10-/f+; vimentin-/+; inhibin-; AMACR-; E-cadherin-/f+; EMA+/- |

| | | | | |
|---|---|---|---|---|
| BHD7 | | Hybrid tumor (with features of clear cell RCC, chromophobe RCC and focally of oncoytoma) | **Growth pattern:** Multilocular, cystic, nested, solid and focally papillary patterns. Highly vascularized connective tissue in between tumor areas and nests. Prominent hemorrhage, frequent foamy macrophages and some hemosiderin macrophages.<br><br>**Cell morphology:**<br><br>Cytoplasm: Mainly pale-reticular and, in areas, clear cytoplasm. Focally strong eosinophilic/ oncocytic cytoplasm. Cell borders are moderately marked except in the oncocytic areas where cell borders are not visible.<br><br>Nucleus: Round / irregular nuclei with prominent central nucleolus. In the oncocytic areas round nuclei without atypia. | CK7-/p+, CK8+/-, EMA+, vimentin-, CD10-, AMCR-/focally+, MelanA-, CA IX- |
| BHD8 | | Hybrid tumor (with features of oncocytoma, chromophobe RCC, and clear cell RCC) | **Growth pattern:** Mainly solid, in places nested in a pale, loose extracellular matrix. Focally also papillary pattern.<br><br>**Cell morphology:**<br><br>Cytoplasm: Mainly oncocytic eosinophilic cytoplasm, in places pale reticular with perinuclear halo. Focally clear cytoplasm. Cell borders vary between invisible and moderately marked.<br><br>Nucleus: Mainly round, focally slightly irregular medium-sized nuclei with clear chromatin. Often central nucleolus that is small or moderately pronounced. Raisinoid nuclei are not a feature. Bi- and multinucleated cell forms are present. | CK7-/f+; CD10+/-; vimentin-; AMACR+/- |

**Table S2.** Methods – Fluorescent cell barcoding. Set-up for 6 conditions for each of the 2 cell lines, resulting in 12 unique populations, as well as 4 compensation controls (1 negative control and 3 single-stained). Fluorophores refer to their respective succinimidyl ester dye as described in Methods, where 10 µl of the annotated dye was used.

<table>
<tr><td>UOK257</td><td>UOK257</td><td>UOK257</td><td>UOK257-2</td><td>UOK257-2</td><td>UOK257-2</td></tr>
<tr><td rowspan="3">30 µL DMSO</td><td rowspan="2">20 µL DMSO</td><td rowspan="2">20 µL DMSO</td><td rowspan="2">20 µL DMSO</td><td>Pacific Blue (5 µg/mL)</td><td>Pacific Blue (5 µg/mL)</td></tr>
<tr><td>10 µL DMSO</td><td>10 µL DMSO</td></tr>
<tr><td>Pacific Orange (2 µg/mL)</td><td>Pacific Orange (80 µg/mL)</td><td>Pacific Blue (5 µg/mL)</td><td>Pacific Orange (2 µg/mL)</td><td>Pacific Orange (80 µg/mL)</td></tr>
<tr><td>UOK257</td><td>UOK257</td><td>UOK257</td><td>UOK257-2</td><td>UOK257-2</td><td>UOK257-2</td></tr>
<tr><td rowspan="2">20 µL DMSO</td><td>10 µL DMSO</td><td>10 µL DMSO</td><td>Pacific Blue (5 µg/mL)</td><td>Pacific Blue (5 µg/mL)</td><td>Pacific Blue (5 µg/mL)</td></tr>
<tr><td>Alexa-488 (5 µg/mL)</td><td>Alexa-488 (5 µg/mL)</td><td>Alexa-488 (5 µg/mL)</td><td>Alexa-488 (5 µg/mL)</td><td>Alexa-488 (5 µg/mL)</td></tr>
<tr><td>Alexa-488 (5 µg/mL)</td><td>Pacific Orange (2 µg/mL)</td><td>Pacific Orange (80 µg/mL)</td><td>10 µL DMSO</td><td>Pacific Orange (2 µg/mL)</td><td>Pacific Orange (80 µg/mL)</td></tr>
<tr><td>Negative control</td><td>Control</td><td>Control</td><td>Control</td><td></td><td></td></tr>
<tr><td>30 µL DMSO</td><td>Pacific Orange (80 µg/mL)</td><td>Pacific Blue (5 µg/mL)</td><td>Alexa-488 (5 µg/mL)</td><td></td><td></td></tr>
</table>

**Table S3.** Alexa Fluor 647 conjugated antibodies used for phospho flow cytometry.

| **Antibody target** | **Provider** | **Catalog number** | **Concentration used** |
|---|---|---|---|
| AKT1 pT308 | CST | 3375 | 1:100 |
| AKT1 pS473 | CST | 4075 | 1:25 |
| BCL2 pS70 | BD | 562531 | 1:20 |
| Histone H3 pS10 | CST | 9716 | 1:67 |
| Isotype control | BD | 557783 | 1:10 |
| MAPK8/JNK pT183/Y185 | CST | 9257 | 1:40 |
| MTOR pS2448 | BD | 564242 | 1:40 |
| STING1 pS366 | CST | 43499 | 1:50 |
| Tyrosine pY | CST | 9415 | 1:50 |
| ULK1 pS757 | CST | 29313 | 1:50 |
| MAPK14/p38 MAPK pT180/Y182 | CST | 4552 | 1:25 |
| MAPK3/p44 and MAPK1/p42 MAPK pT202/Y204 | CST | 4375 | 1:100 |
| RPS6 pS235/236 | CST | 4851 | 1:100 |
| TBK1 pS172 | BD | 558603 | 1:40 |
